# Functional genomics identifies extension of complex N-glycans as a mechanism to evade lysis by natural killer cells

**DOI:** 10.1101/2023.04.03.535404

**Authors:** Xiaoxuan Zhuang, James Woods, Yanlong Ji, Sebastian Scheich, Fei Mo, Matthias Voss, Henning Urlaub, Kuan-Ting Pan, Eric O. Long

## Abstract

Somatic mutations can lead to the transformation of healthy cells into malignant cells and allow their evasion from immune surveillance. To uncover genes that play a role in the detection and lysis of tumor cells by natural killer (NK) cells, a B lymphoblastoid cell line was subjected to a genome-wide CRISPR screen. Among the top hits that facilitated NK evasion was *SPPL3*, which encodes an intramembrane protease that cleaves transmembrane glycosyltransferases in the Golgi apparatus. *SPPL3*-deficient cells accumulated glycosyltransferases, such as acetylglucosaminyltransferase 5 (MGAT5), and displayed increased N-glycosylation. Binding of NK receptors NKG2D and CD2 to their corresponding ligands MICB and CD58, and binding of rituximab to CD20, was disrupted by *SPPL3*-deletion. Inhibition of N-glycan maturation restored receptor binding and sensitivity to NK cells. To elucidate the mechanism of this resistant phenotype, a secondary CRISPR screen was performed in *SPPL3*-deficient cells. This screen identified glycosyltransferases that catalyze the formation of highly branched N-glycans and N-acetyl-lactosamine (LacNAc) extensions as key regulators that prevent killing. A significant enrichment of poly-LacNAc-containing tetra-antennary species was confirmed by glycoproteomic analysis. These findings provide mechanistic insight into how *SPPL3* deletions have been linked to cancer.

## Introduction

Natural Killer (NK) cells play a crucial role in the immune system due to their ability to recognize and kill infected or abnormal cells, including cancer cells [1]. NK cells detect target cells through a combination of activating and inhibitory receptors. Activating receptors, such as the natural cytotoxicity receptors (NCRs), NKG2D, and 2B4, recognize stress-induced ligands on the surface of target cells. Inhibitory receptors, such as killer cell immunoglobulin-like receptors (KIR) and NKG2A, bind to class I human leukocyte antigens (HLA-I) on healthy cells [1–4]. If the balance of activating and inhibitory signals favors activation, NK cells form tight conjugates with target cells and release the content of lytic granules to induce apoptosis in the target cell [1]. Alternatively, NK cells expressing TRAIL (TNFSF10) or Fas ligand (FASLG) engage death receptors on target cells, such as TNFRSF10 or FAS, to trigger apoptotic death. A positive correlation between the presence of NK cells and the response to immunotherapy in cancer patients has been established [5, 6]. NK-focused cancer immunotherapies have shown promise for cancer treatment [7, 8]. Due to a milder cytokine response and lower risk of graft versus host disease (GvHD) than T cells in allogeneic settings, NK cells are ideal candidates for use as effector cells to express chimeric antigen receptors (CAR) for immunotherapy, particularly in the development of “off-the-shelf” products [9]. Besides the killing induced by CARs, NK cell activating receptors engage with ligands on tumor cells and trigger additional lysis, which can provide patients with extra beneficial effects from NK-based cell therapy. The first in-human phase 1 and 2 trials using NK cells carrying a CD19-CAR to treat patients with CD19-positive non-Hodgkin’s lymphoma and chronic lymphoblastic leukemia have proven safe and showed encouraging responses [8].

Given the promise of NK cells as effector cells for use in treating certain cancers, it is important to determine how cancer cells could evade recognition and killing by NK cells. Although much is known about receptor-ligand interactions that regulate target cell lysis by NK cells, other pathways that may render cancer cells resistant or sensitive to NK cells have not been fully explored. These could include various intracellular changes in metabolism, lipid composition, membrane organization, and many other factors. To discover these resistance and sensitivity factors, we carried out CRISPR screens in a chronic myeloid leukemia (CML) cell line and uncovered the IFN-γ signaling pathway as a strong contributor to tumor evasion [10], as confirmed in recent studies [11–13]. By upregulating expression of HLA-I molecules on target cells, IFN-γ signaling increases the engagement of inhibitory receptors on NK cells. Conversely, the same pathway has the opposite effect on recognition and killing of tumor cells by cytotoxic CD8 T cells, due to increased presentation of tumor antigen by HLA-I [14]. The sensitivity to NK cells varies widely by tumors and cell types and is likely regulated through different pathways.

B cell malignancies, such as chronic lymphocytic leukemia (CLL) and non-Hodgkin lymphoma (NHL), are cancers that originate from B lymphocytes. These cancers cells often evade the immune system, including their detection by NK cells, due to various mechanisms such as downregulation of cell surface ligands that are essential for NK cell recognition and activation. For instance, mutations or deletions in *CD58*, which encodes the ligand for human receptor CD2, occur in approximately 21% of diffuse large B-cell lymphomas (DLBCL) [15]. Despite these evasion mechanisms, high NK cell count remains a valid predictor of response to immunotherapy for patients with B cell malignancies [16].

Therefore, to gain a deeper understanding of the interaction between NK cells and B-cell malignancies, we performed here a genome-wide CRISPR knockout screen in the 721.221 cell line (referred to as 221), an EBV-transformed lymphoblastoid B-cell line. This cell line had been selected for the loss of classical HLA-I genes [17] and is therefore sensitive to lysis by NK cells. As expected, the loss of ligands for NK activation receptors resulted in reduced sensitivity. Deletion of several other genes, including signal peptide peptidase-like 3 (*SPPL3)*, had the greatest impact on resistance to killing by NK cells. Here, we describe how *SPPL3* functions as a crucial controller of the interaction between NK cells and their targets. We found that the increased presence of complex N-glycans on the surface of *SPPL3*-deleted tumor cells limits the binding of NK cell receptors and facilitates immune evasion. We further identified glycosyltransferases that were responsible for this resistance phenotype by performing a secondary CRISPR screen in *SPPL3*-deleted cells.

## Results

### CRISPR-Cas9 screen to identify genes that regulate sensitivity to killing by NK cells

221 cells, first stably transfected with Cas9, were then stably transduced with the GeCKO V2 CRISPR library. After 10 days of puromycin selection, 221 cells were co-cultured with IL-2-expanded NK cells at low effector to target ratios (E:T of 1:10 to 1:5) until about 10% to 20% of 221 cells had survived. Surviving 221 cells were harvested for sequencing (Fig. 1A) and a lysis assay was performed to confirm reduced sensitivity (Fig. 1B). Surviving 221 cells were more resistant to NK-mediated lysis compared to 221 cells that had been transduced with the GeCKO V2 CRISPR library but not selected by co-culture with NK cells (Fig. 1B). Analysis of both enriched and depleted guide RNAs (sgRNAs) identified novel genes that regulate sensitivity or resistance to NK-mediated killing (Fig. 1C, D). The top 30 enriched genes included *CD58*, a known NK activating ligand, and other regulators such as *FBXO11* and *SPPL3* (Fig. 1C). Conversely, the top 30 depleted genes included *B2M* and other resistance-regulators such as *EP300* and *ADAM2* (Fig. 1D). Gene set enrichment analysis revealed that sgRNAs targeting genes in the IFN-γ and antigen presentation pathways were depleted (Fig. 1E), while sgRNAs targeting other ligands for NK cell receptors such as MICB, PVR, TNFRSF10B and TNFRSF10A were enriched (Fig. 1F). These ligands engage NK receptors NKG2D, DNAM-1 (CD226), and TNFSF10 (TRAIL), respectively.

**Fig. 1.**
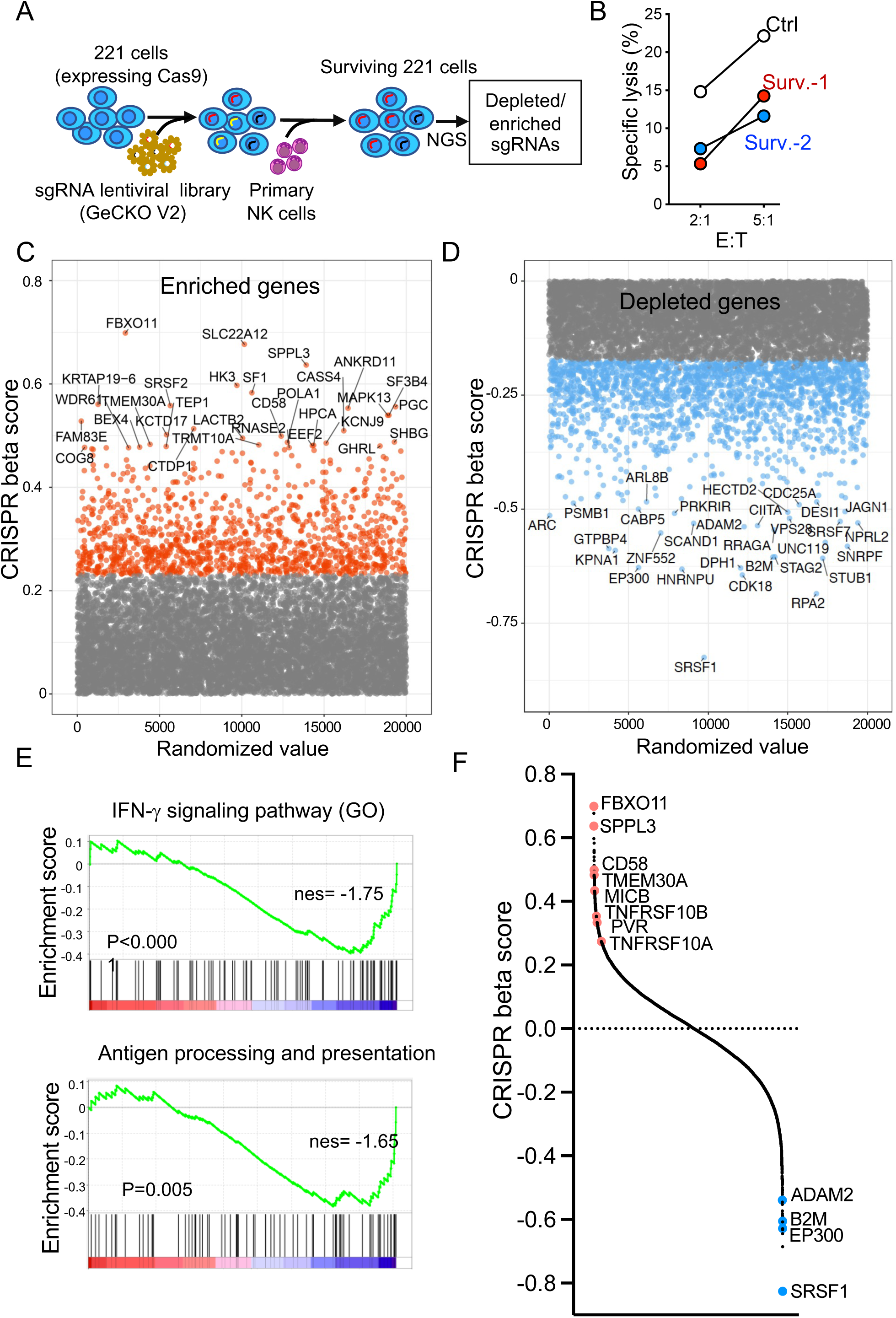
CRISPR-Cas9 screen to identify genes that regulate sensitivity to NK cells. (A) Experimental outline of the genome-wide CRISPR knock out screen in 221 cells. 221-Cas9 cells transduced with the sgRNA library were co-incubated with activated primary human NK cells. Distribution of sgRNAs in surviving 221 cells were analyzed by NGS. (B) Representative killing assay by flow cytometry with 221 cells that survived co-culture with NK cells (Surv.-1 and Surv.-2) or 221 cells cultured without NK cells (Ctrl). (C-D) Scatter plots showing enriched (C) and depleted (D) genes in the screen. CRISPR scores were calculated via the MAGeCK MLE analysis (y axis). Each dot represents a gene, and the top 30 enriched (C) and depleted (D) genes are labeled. (E) GSEA analysis of CRISPR scores of depleted genes and their relationship with IFN-γ signaling and antigen processing and presentation pathways. (F) Ranked plot of CRISPR scores from the screen in 221 cells. Highlighted genes are color-coded red for enriched genes and blue for depleted genes.

To validate the top hits, we performed killing assays on 221 cells edited with non-targeting sgRNA (sgNT) or sgRNA targeting a single gene of interest. These cells were pre-labeled with different combinations of fluorescent membrane dyes (Fig. 2A, B), mixed with NK cells at a 1:1 ratio and incubated for 16 hours with NK cells (Fig. 2A). Flow cytometry was used to determine changes in cell ratios after co-culture. Knockouts of NK receptor ligands MICB, CD58, TNFRSF10B and TNFRSF10A were more resistant to NK cell-mediated killing compared to sgNT 221 cells (Fig. 2B). The knock-out efficiency of sgRNAs targeting these genes was validated by staining with respective antibodies (Fig. 2C). Deletions of regulators *FBXO11* and *SPPL3* resulted in greatly reduced sensitivity to killing (Fig. 2B), and the sgRNAs targeting these genes efficiently reduced their expression (Fig. 2D and E). 221 cells expressing sgRNA for *MICB*, a ligand of receptor NKG2D, stimulated less degranulation by NK cells (Fig 2F). Deletion of *CD58*, the ligand for CD2, had a greater impact on killing but did not change NK degranulation by NK cells (Fig. 2F). This is consistent with the function of CD2 in adhesion and of other activation receptors on degranulation [18–20]. Knockouts of *SPPL3* and *FBXO11* also resulted in a reduced stimulation of NK cell degranulation (Fig. 2G).

**Fig. 2.**
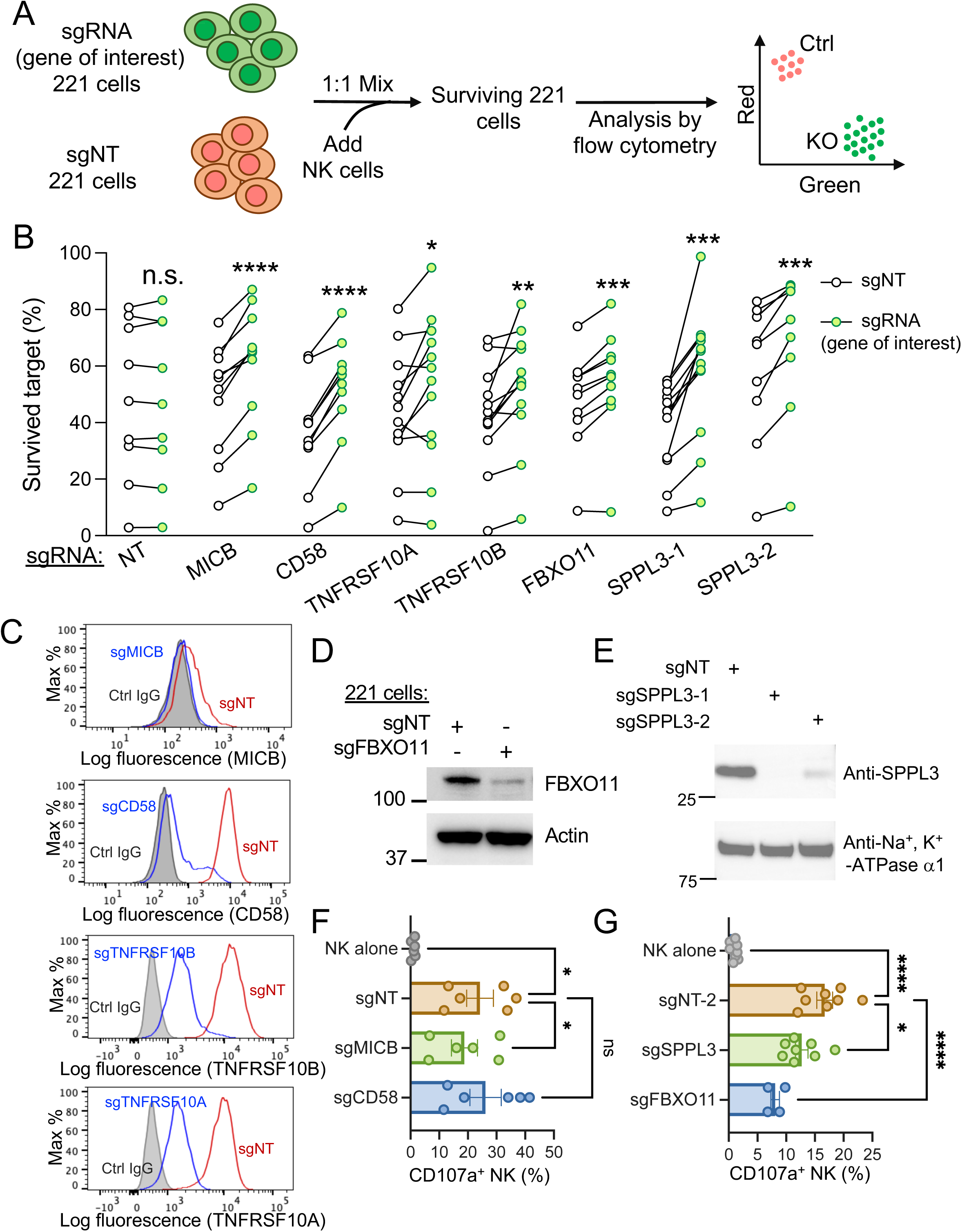
Validation of CRISPR screen hits. (A) Experimental outline of the killing assay used to validate hits from the CRISPR screen. 221 cells expressing non-targeting sgRNA (sgNT) and sgRNA targeting genes of interest were prelabeled with dyes and mixed at a 1:1 ratio before co-culture with NK cells. Surviving 221 cells were analyzed by flow cytometry. (B) Survival of 221 target cells expressing sgRNAs for the indicated genes (green dots) compared to cells expressing sgNT (white dots). *SPPL3* was knocked-out separately with two different sgRNAs. Each line represents an independent experiment with NK cells from different donors (n≥ 8, paired t-test, n.s. not significant, *P<0.05, **P<0.01, ***P<0.001, ****P<0.0001). (C) Representative histograms showing staining of the indicated ligands of NK cell receptors on 221 cells expressing sgNT or sgRNAs targeting the respective ligands. (D) Immunoblot of FBXO11 from a lysate of 221 cells expressing sgNT or sgFBXO11. Immunoblot of actin was used as loading control. (E) Immunoblot of SPPL3 isolated from the purified membrane fraction of 221 cells that expressed sgNT or sgSPPL3. Immunoblot of the Na^+^, K^+^-ATPase was used as loading control. (F) NK cell degranulation induced by 221 cells expressing sgNT or sgRNAs targeting genes encoding the NK cell ligands MICB and CD58. Data shown as mean ± SEM (n=6, *P<0.05, n.s. not significant). (G) NK cell degranulation induced by 221 cells expressing sgNT or sgRNAs targeting SPPL3 and FBXO11. Data shown as mean ± SEM (n=8, one-way ANOVA test, *P<0.05, ****P<0.0001).

### Increased glycosylation of ligands for NK receptors and reduced binding of soluble receptors to *SPPL3*-deleted cells

To further investigate the effect of SPPL3 deletion, we examined the expression of several ligands of NK receptors on SPPL3-deleted 221 cells (Fig. S1A-E). Surface levels of CD48, CD58, ICAM-1, CD80, HLA-E, and MICB were unchanged in SPPL3-deficient cells (Fig. S1A and B). RNA-seq analysis was performed to examine the expression of various ligands of NK receptors in sgNT and sgSPPL3 221 cells. The cells were clustered into respective groups in a Principal Component Analysis (PCA) (Fig. S1C). The transcriptional levels of ligands such as TNFRSF1A/B, TNFRSF10A/B, SLAMF7/6, PVR, MICB, ICAM1, HLA-E, FAS, CD80, CD70, CD58, CD48, and CD27 were not affected (Fig. S1D). SPPL3 deletion resulted in minor changes in the transcriptome of 221 cells, with a significant upregulation of only 2 genes, and downregulation of 6 genes (Fig. S1E).

SPPL3 is a Golgi-resident intramembrane protease that cleaves the catalytic domain of glycosyltransferases from their membrane anchors, which could promote their secretion into the extracellular space and lead to a reduced intracellular abundance and thus activity of these enzymes [18, 21]. MGAT5 is one of the best-studied glycosyltransferases cleaved and regulated by SPPL3 [21]. We monitored MGAT5 protein levels in cell lysates and supernatants of 221 cell cultures (Fig. 3A). SPPL3-deficient 221 cells had increased cellular expression of MGAT5 and a decreased amount of MGAT5 in the culture media, suggesting that MGAT5 is retained inside SPPL3-deficient cells where it promotes increased glycosylation and branching of N-glycans (Fig. 3A). To test this, we stained 221 cells with the lectin Con A to detect high mannose and SNA to determine the extent of sialylation. SPPL3 knockout cells had reduced high mannose and increased terminal sialic acid glycans (Fig. 3B), indicating a shift towards more complex glycans. To examine the glycosylation status of several ligands for NK receptors, we performed motility shift experiments and found that glycosylation of CD58 was greatly increased in SPPL3 knockout 221 cells (Fig. 3C). Similarly, HA-tagged versions of MICB and PVR transfected into 221 cells showed reduced mobility in SPPL3 knockout cells (Fig. 3D). To test whether increased glycosylation of ligands could interfere with receptor interaction, we measured binding of soluble, recombinant receptors NKG2D and CD2 to sgSPPL3 221 cells. Binding of either one to SPPL3-deficient cells was decreased, as compared to binding with control sgNT 221 cells (Fig. 3E and F), presumably due to steric hindrance by the larger N-glycans on MICB and CD58.

**Fig. 3.**
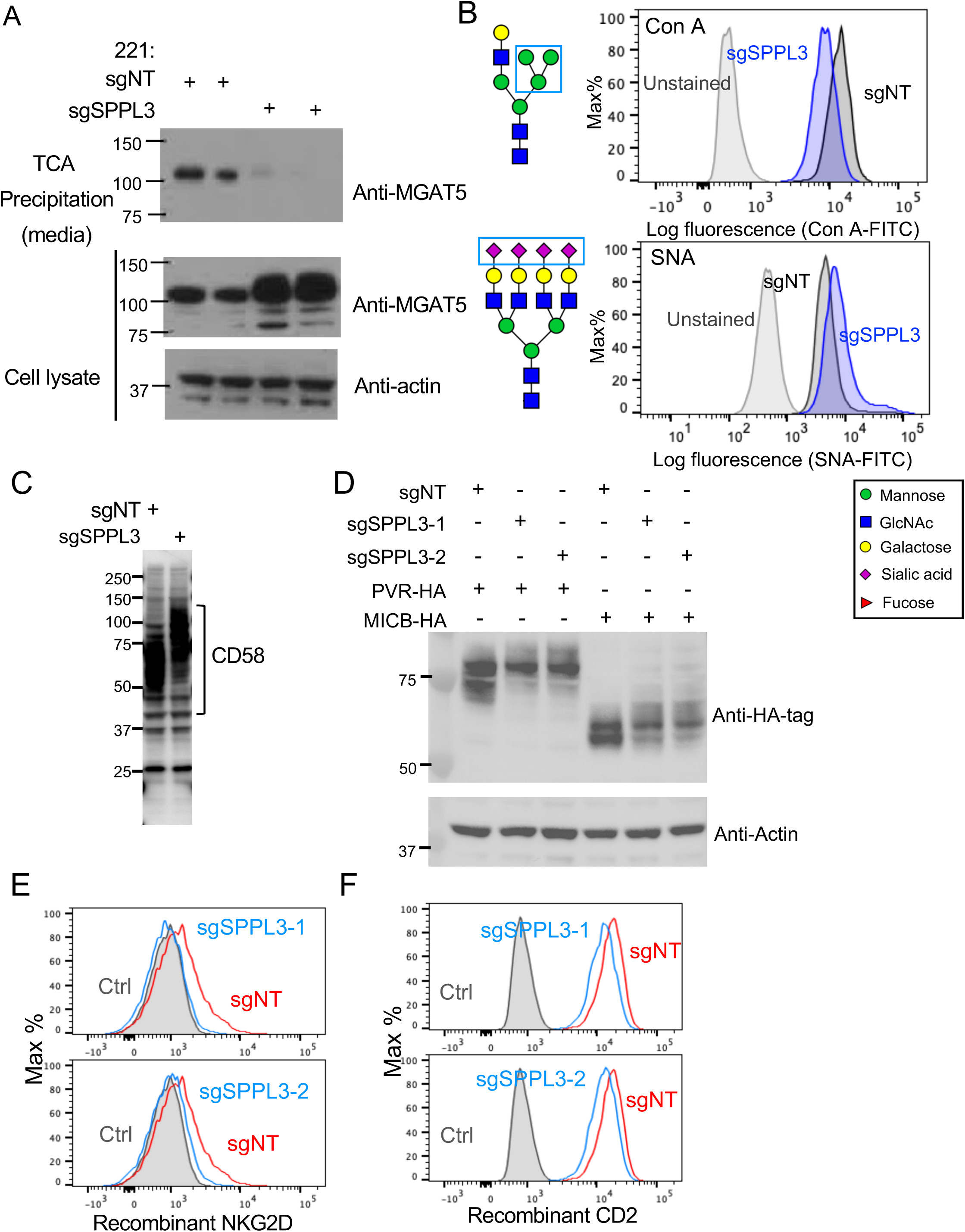
Impact of SPPL3 deletion on glycosylation of ligands for NK receptors and on binding to their receptors. (A) Immunoblot of MGAT5 in TCA-precipitated culture supernatants (top panel) and in total cell lysates (center panel) of sgNT-expressing and SPPL3 knockout cells. Immunoblot of actin was used as loading control (bottom panel). (B) Representative histograms of lectins Con A and SNA binding to 221 cells expressing sgNT or sgSPPL3 (blue). The binding specificities for glycan structures (blue box) are shown using the standard symbols for glycans (shown in box). (C) Immunoblot of CD58 in total cell lysates of SPPL3 knockout 221 cells versus sgNT 221 cells. (D) Immunoblot of HA-tagged PVR and MICB transfected into SPPL3-deleted 221 cells and sgNT 221 cells. Whole cell lysate was immunoblotted with antibodies to HA and to actin. (E) Representative histograms show binding of soluble recombinant NK cell receptors NKG2D and CD2 to 221 cells transfected with sgNT (red) or sgSPPL3 (blue).

If our interpretation was correct, the effect of *SPPL3* deletion and enhanced N-glycosylation should be reversed by inhibition of the N-glycan pathway. Therefore, we treated both sgNT and sgSPPL3 221 cells with kifunensine, a potent inhibitor of mannosidase I (Fig. 4A). Trimming of high mannose forms by mannosidase I is essential for the synthesis of complex N-glycan forms. Using the lectin ConA specific for terminal mannose residues, which are exposed prior to trimming, we observed much higher staining of kifunensine-treated sgNT and sgSPPL3 cells, as compared to untreated cells (Fig. 4A, B). After treatment with kifunensine, the difference in ConA staining between untreated sgNT cells and sgSPPL3 cells was eliminated (Fig. 4B). Kifunensine also prevented the maturation of large glycans on CD58 and MICB, as determined by electrophoretic motility (Fig. 4C and D). Notably, kifunensine treatment restored NK cell degranulation stimulated by SPPL3-deficient 221 cells (Fig. 4E) and reinstated the binding of soluble recombinant CD2 to these cells (Fig. 4F).

**Fig. 4.**
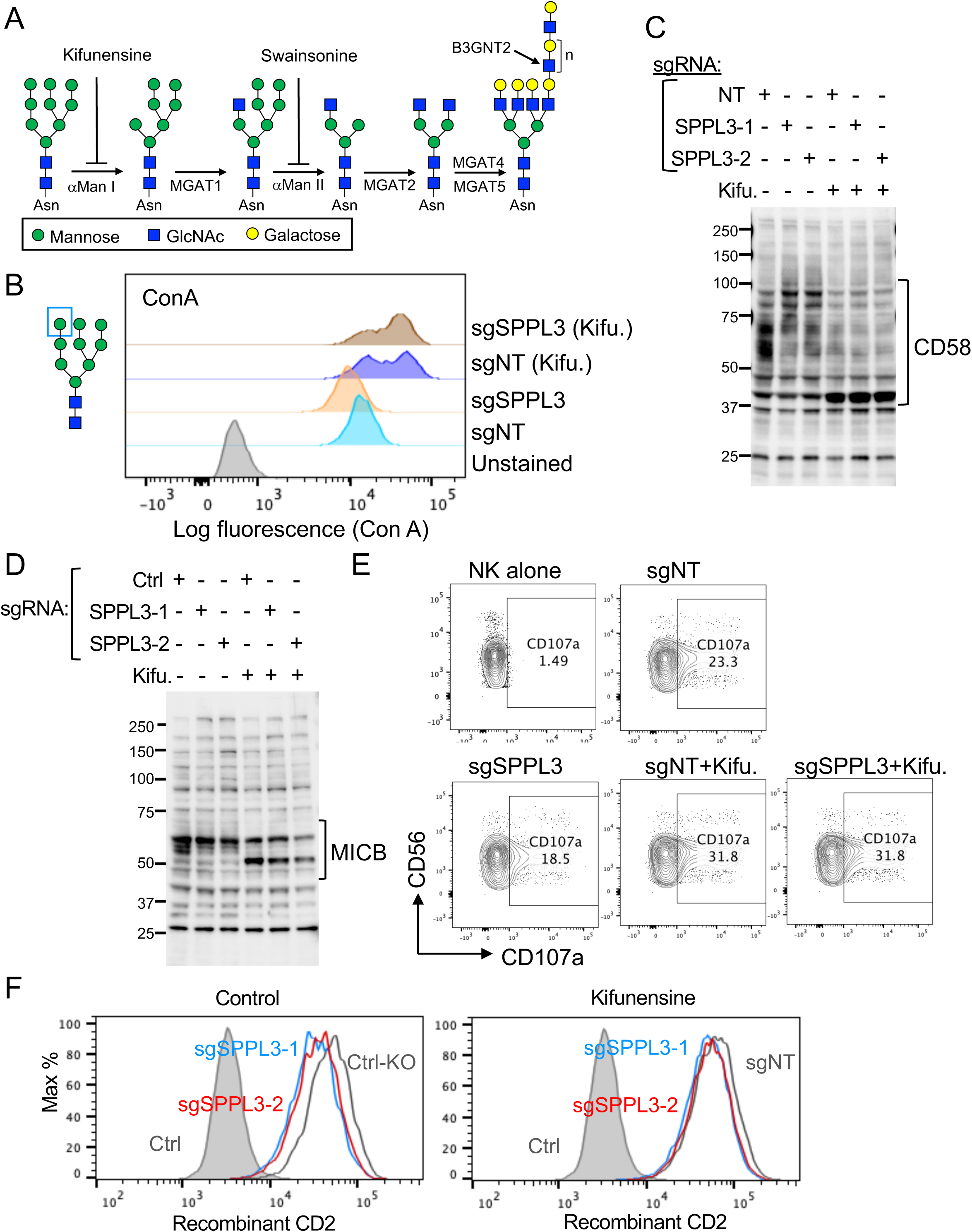
Inhibition of N-glycan maturation reduced the glycosylation of ligands for NK receptors, and restored NK cell degranulation induced by SPPL3 knockout 221 cells. (A) Diagram of N-glycan maturation from the high-mannose stage. The mannosidase I inhibitor kifunensine and the mannosidase II inhibitor swainsonine inhibit the trimming of high mannose glycans. (B) Con A staining of CD58 in sgNT and sgSPPL3 221 cells, treated or not with kifunensine, as indicated. (C-D) Immunoblots of CD58 (C) and MICB (D) in sgNT and SPPL3 knockout 221 cells, treated or not with kifunensine. (E) Representative contour plots of surface CD107a expression on NK cells stimulated with sgNT and sgSPPL3 221 cells. (F) Representative histograms of recombinant CD2 binding to sgNT and sgSPPL3 cells before and after kifunensine treatment.

### Disruption of the N-glycan maturation pathway increased the sensitivity of *SPPL3*-deleted 221 cells to NK-mediated killing

The sensitivity of kifunensine-treated 221 cells to NK-mediated killing was evaluated next. sgNT and sgSPPL3 221 cells treated or not with kifunensine were barcoded using different concentrations of PHK26 (red) and PKH67 (green) dyes and mixed at a 1:1:1:1 ratio (Fig. 5A). Their relative sensitivity to NK-mediated killing was then assessed using flow cytometry (Fig. 5A). SPPL3-deleted cells were more resistant to killing than sgNT cells, and kifunensine treatment reverted the sensitivity of SPPL3-deleted cells back to the level of killing observed with sgNT-treated 221 cells (Fig. 5B). To confirm the specificity of kifunensine, we repeated the killing assay using another glycosylation inhibitor, swainsonine, which blocks a later step in glycan maturation executed by the Golgi-resident alpha-mannosidase II (Fig. 4A). Similarly, the greater resistance of sgSPPL3 cells to NK-mediated killing disappeared after swainsonine treatment (Fig. 5C). Both kifunensine and swainsonine reduced the amount of complex glycans at the cell surface that were detected by the lectin PHA-E on sgNT and sgSPPL3 221 cells (Fig. S2).

**Fig. 5.**
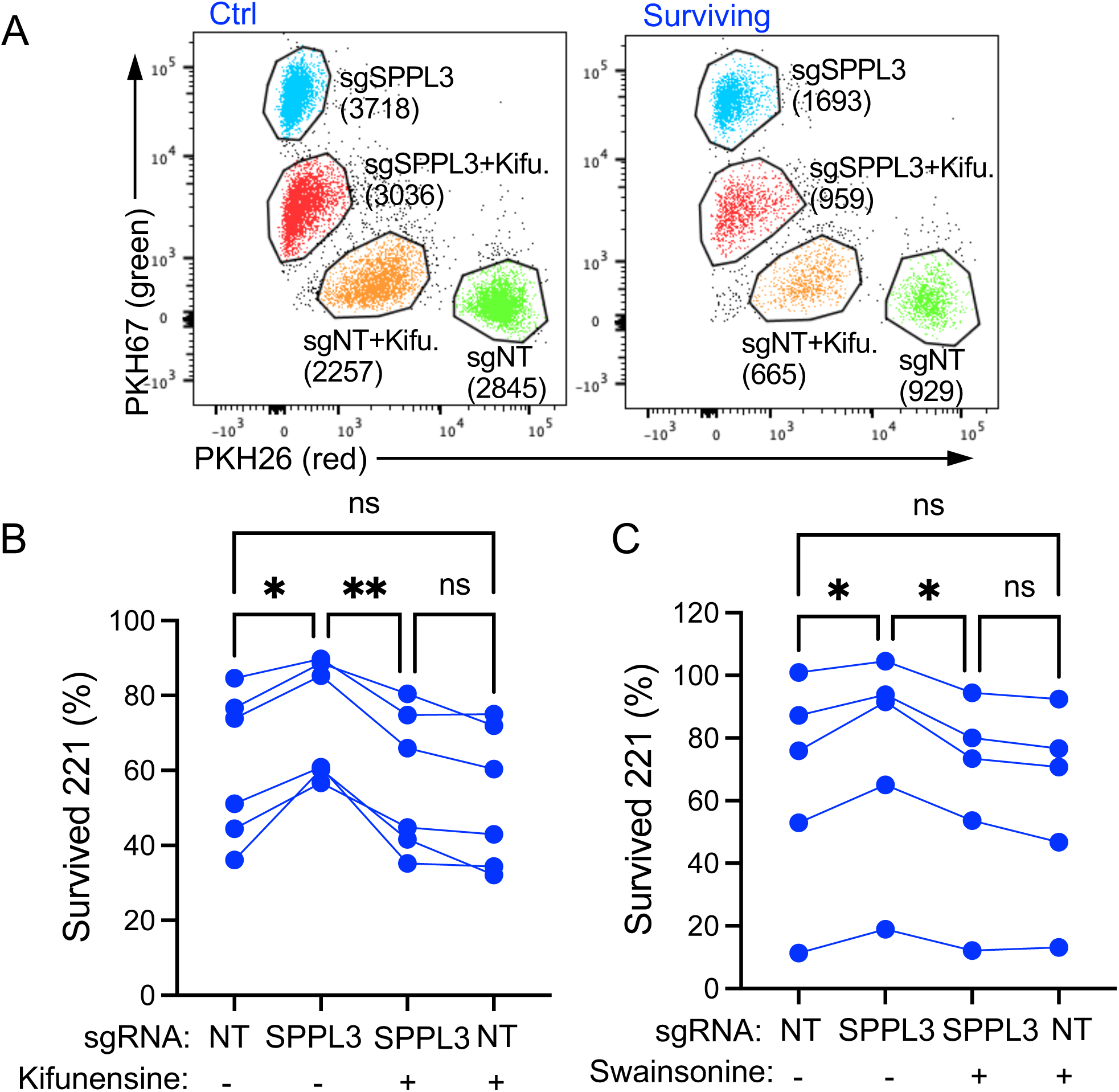
Treatment with N-glycosylation inhibitors restored the sensitivity of SPPL3-deleted 221 cells to NK-mediated killing. (A) Representative dot plots of flow cytometry analysis of color-coded sgNT or sgSPPL3 221 target cells in a killing assay with NK cells. Cells were treated or not with Kifunensine. (B) Statistics of independent donors as in (A). Each line represents a different donor (n=6, paired one-way ANOVA test, n.s. not significant, *P<0.05, **P<0.001). (C) Statistics of color-coded killing assays to compare sensitivity of sgNT and sgSPPL3 expressing 221 cells, which were either non-treated or treated with the Golgi alpha-mannosidase II inhibitor swainsonine. Each line represents an independent donor (n=6, paired one-way ANOVA test, n.s. not significant, *P<0.05)

In addition to impairment of receptor-ligand interactions at the NK-target cell synapse, heavy glycosylation may also impair the binding of antibodies to tumor cell antigens and prevent the ability of NK cells to perform antibody-dependent cellular cytotoxicity (ADCC). This is relevant in the context of monoclonal antibody therapies that target tumor antigens. The CD20 antibody rituximab is one of the most established antibodies used to treat various B-cell malignancies [22]. We observed that binding of rituximab to 221 cells deficient in SPPL3 was indeed reduced (Fig. 6A and B). Cells lacking SPPL3 were more resilient to CD20-dependent NK cell-mediated ADCC, and their resistance to ADCC was restored by treatment with kifunensine (Fig. 6C).

**Fig. 6.**
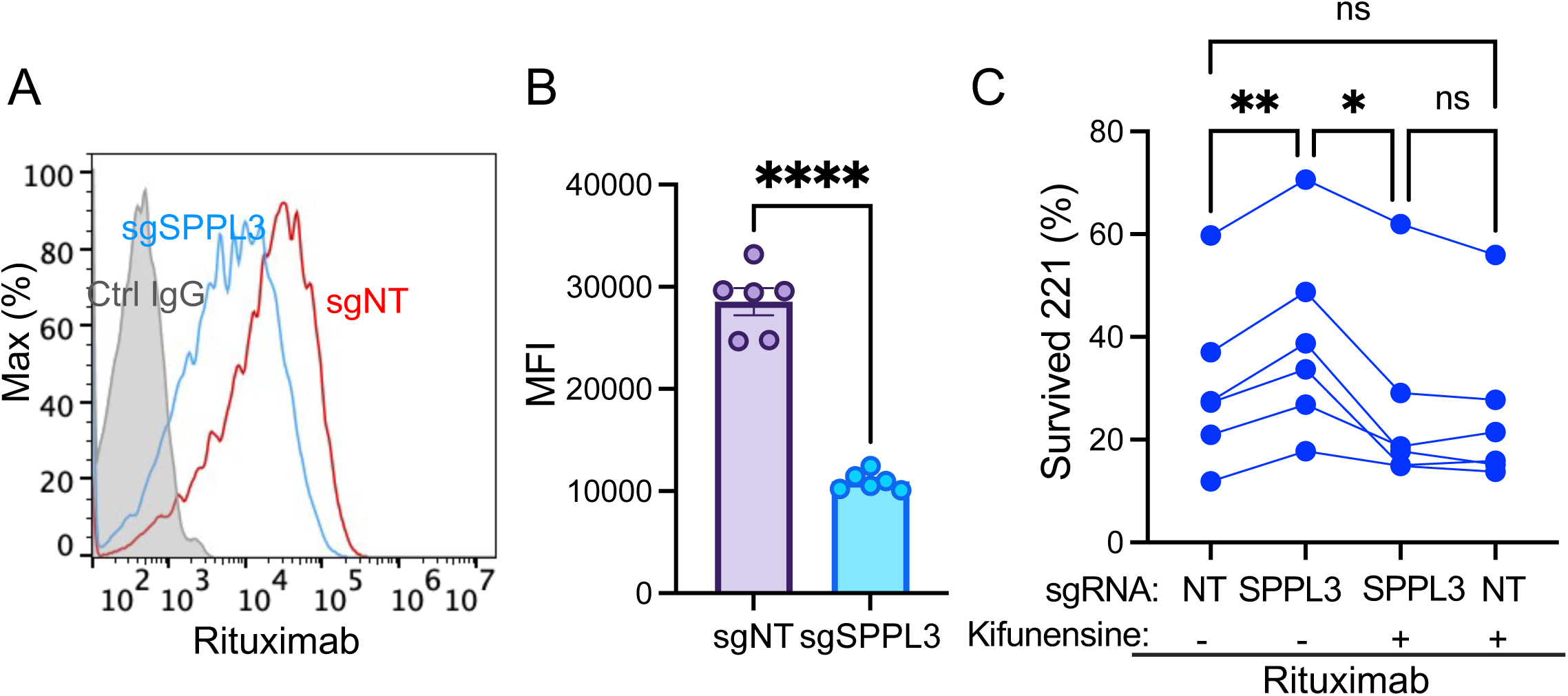
SPPL3 deletion in 221 cells reduces rituximab binding to CD20 and decreases NK-mediated ADCC. (A) Histograms of rituximab staining of 221 cells expressing sgNT (red) or sgSPPL3 (blue). (B) Statistics of several experiments performed as in (A). Data is shown as mean ± SEM (n=6, unpaired t-test, ****P<0.0001). (C) Statistics from killing assays to compare the sensitivity of 221 cells expressing sgNT or sgSPPL3 to NK-mediated ADCC triggered by 10 μg/mL rituximab. Cells were treated or not with Kifunensine, as indicated. Each line represents NK cells from different donors (n=6, paired one-way ANOVA test, n.s. not significant, *P<0.05, **P<0.01)

### A secondary CRISPR screen in *SPPL3*-knock out cells to identify SPPL3 substrates that are responsible for resistance to NK cell-mediated lysis

To gain insight into how glycan structures of ligands on 221 cells for NK receptors can impact the sensitivity of 221 cells to NK-dependent killing, a CRISPR screen focused on glycosylation pathways [23] was performed with *SPPL3*-knockout cells (Fig. 7A). Cumulative CRISPR scores (CSS) were determined by subtracting the CSS from sgNT cells from the CSS from sgSPPL3 cells, as displayed in Fig. 8B. *B3GNT2* was one of the top hits of deleted genes that restored sensitivity of *SPPL3*-knockout cells to NK-dependent killing (Fig. 7A and B). *B3GNT2* encodes the main glycosyltransferase that extends highly branched N-glycans by transfer of a GlcNAc moiety in a β1,3 linkage with a terminal unsialylated Galactose, preferentially on the MGAT5-generated β-1, 6-linked branch in tri- and tetra-antennary N-glycans [24]. The subsequent transfer of Gal in a β1,4 linkage to GlcNAc by a B4GALT transferase generates the di-saccharide N-acetyl-lactosamine (LacNAc). Iterations of B3GNT2 and B4GALT-mediated transfers generate additional poly-LacNAc extensions (Fig. 4A). We used the lectin LEL, which binds preferentially to LacNAc, to test for the presence of LacNAc on 221 cells. The stronger binding of LEL to the plasma membrane of sgSPPL3 cells, as compared to sgNT 221 cells, indicated an increased amount of LacNAc (Fig. 7C). Deletion of *B3GNT2* in both sgNT and sgSPPL3 cells drastically reduced LEL binding to these cells (Fig. 7D). In addition, the loss of B3GNT2 in 221 cells restored the ability of NK cells to degranulate in response to stimulation by *SPPL3*-knockout cells (Fig. 7E). These findings suggest that the reduced activation of NK cells by *SPPL3*-deleted 221 cells is due to the presence of complex N-glycan structures, including tri- and tetra-antennary branching and their extension with additional LacNAc.

**Fig. 7.**
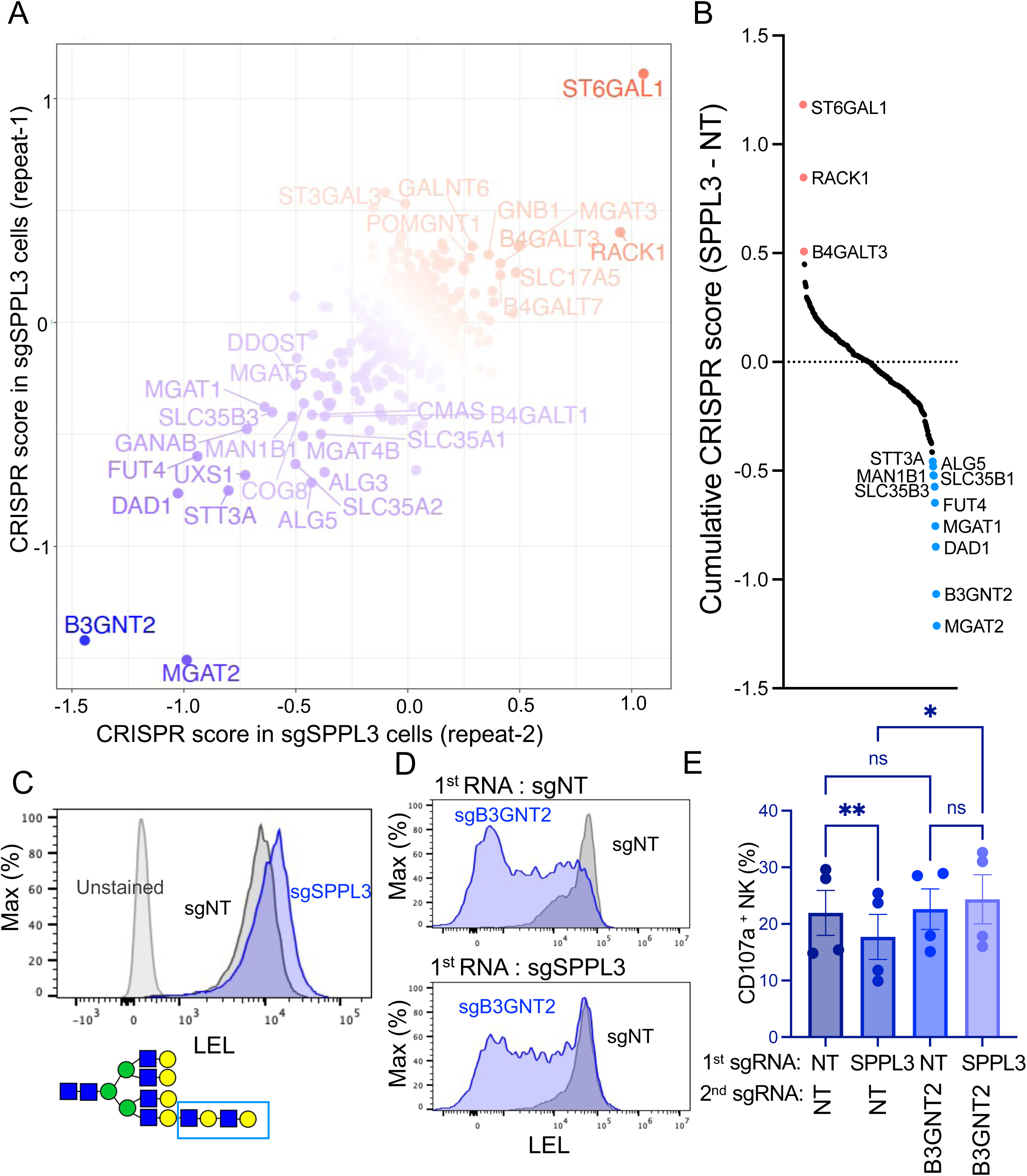
Secondary CRISPR screen in SPPL3-deficient 221 cells to identify glycosyl transferases responsible for the inhibition of NK-mediated killing. (A) CRISPR knockout screen in *SPPL3*-deleted 221 cells using a glycosylation-focused sgRNA sub-library. 221 cells expressing the library were co-cultured with activated primary NK cells. Distribution of sgRNAs in surviving 221 cells was analyzed by NGS. Two biological repeats were plotted on the x-axis and y-axis, respectively. CRISPR beta scores were calculated using the MAGeCK MLE method. Enriched genes, signifying greater resistance, are shown in red, whereas depleted genes, signifying greater sensitivity, are in blue. Gene names of top hits are shown. (B) Ranked plot of cumulative CRISPR scores calculated by the score from SPPL3 knockout cells minus the score from 221 cells expressing sgNT. (C) Representative histograms of LEL binding to sgNT (top) or sgSPPL3 (lower) 221 cells. (D) Representative histograms of LEL binding to SPPL3-KO 221 cells expressing two individual sgRNAs with the 1^st^ sgRNA being sgNT (upper) or sgSPPL3 (lower) and the 2^nd^ sgRNA either non-targeting or against B3GNT2. (E) NK cell degranulation stimulated by 221 cells expressing two individual sgRNAs as in (D). Each dot represents NK cells from an independent donor (n=4, paired one-way ANOVA test, ns not significant, *P<0.05, **P<0.01).

**Fig. 8.**
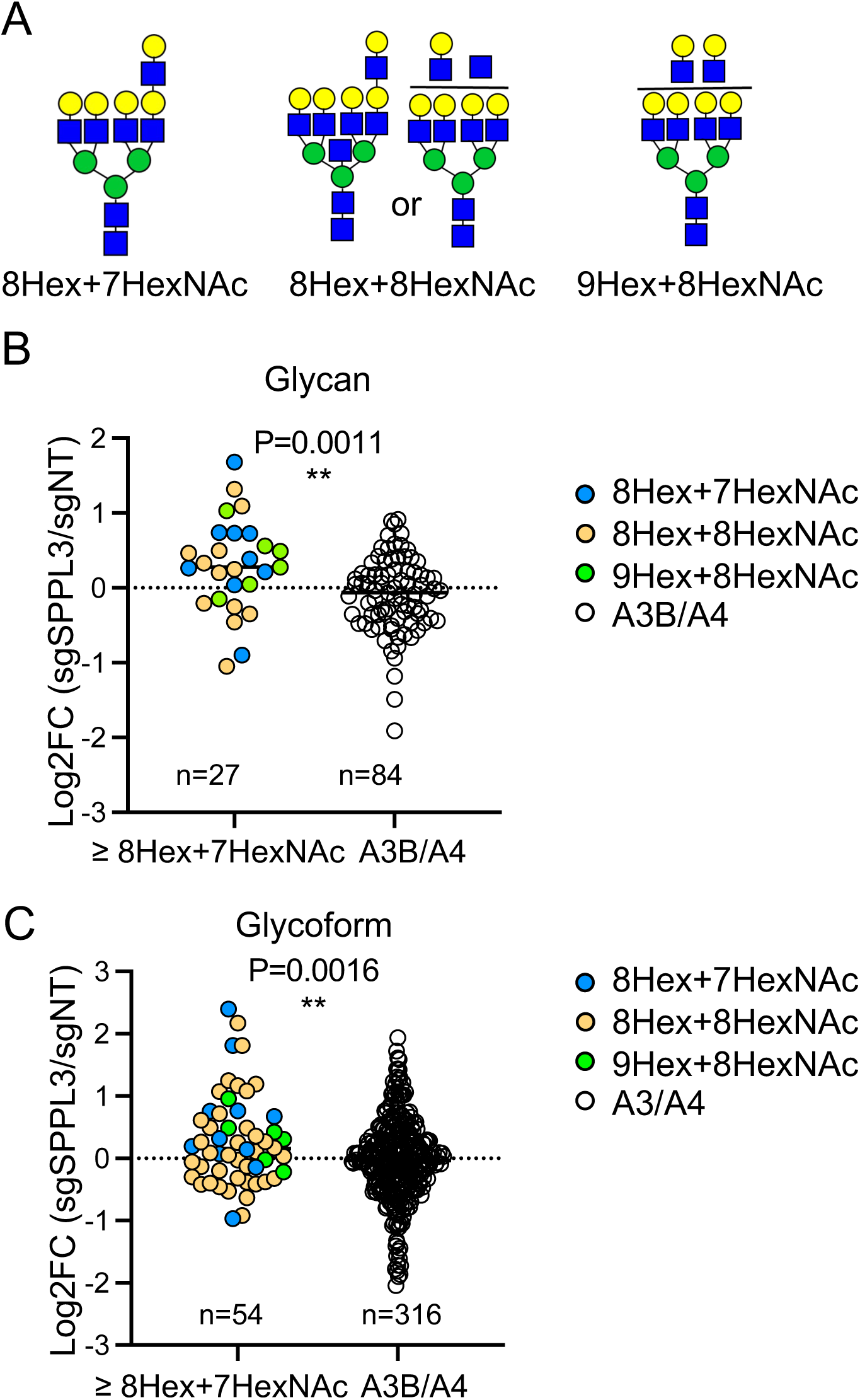
High molecular mass N-glycans on SPPL3-deficient cells. (A) Diagram of N-glycans carrying at least 8 hexoses (Man and Gal) and 7 HexNacs (GlcNAc) that were identified by glycoproteomic analysis of sgNT and sgSPPL3 221 cells. (B) Relative abundance of glycans shown in (A) compared to all other tri- and tetra-antennary (A3B/A4) glycans. (C) Same comparison as in (B) but for glycoforms. Log2FC of relative abundance in SPPL3-deleted cells compared to sgNT cells is shown on the y-axis (unpaired t-test, **P<0.01).

Furthermore, the screen showed that deletion of other genes involved in N-glycosylation, such as *MGAT2* and *MGAT1*, and *FUT4*, resulted in increased sensitivity of *SPPL3*-knockout cells (Fig. 7B), indicating that branching by MGATs and fucosylation by FUT4 of complex N-glycans contributed to resistance.

### Glycoproteomic and glycomic analysis of N-glycans on cells lacking SPPL3

Analysis of the glycoproteome of sgSPPL3 and sgNT 221 cells revealed 273 unique glycans and 5132 glycoforms. Fig. 8A displays the structure of identified N-glycans that carried 4 branches and elongation of at least one branch. Cells lacking SPPL3 had a higher abundance of such complex N-glycans (having at least 8 Hex and 7 HexNAc), as compared to the control cells (Fig. 8B). This trend was confirmed by analyzing unique glycoforms, which showed an increase in complex glycan structures in the *SPPL3*-knock out cells (Fig. 8C). The glycoproteome analysis identified high molecular mass glycans on abundant proteins only. ICAM-1, a ligand for adhesion of NK cells, and CD48 each carried at least one tetra-antennary glycan that was elongated with terminal LacNAc moieties (Table S1). To obtain more information on the composition of N-glycans in *SPPL3*-knock out cells, we performed a glycomic analysis with a focus on larger N-glycan species (Fig. S3). The *SPPL3* knockout resulted in an increase of complex N-glycans, including higher mass glycans carrying poly-LacNAc (Fig. S3).

## Discussion

The goal of this study was to identify genes that promote resistance of target cells against NK cell attack. Such information could point to mutations that may confer resistance to cancer cells. One of the genes with the strongest evasion scores in a genome-wide CRISPR screen for resistance to NK cells was *SPPL3*. SPPL3 is an intramembrane protease located in the Golgi apparatus that cleaves glycosyltransferases in their transmembrane region and releases their catalytic domain [18, 21, 25]. The activity of SPPL3 reduces the accumulation of intracellular glycosyl-transferases and the extent to which proteins are glycosylated during transport through the Golgi. Accordingly, *SPPL3* deletion in the B-cell lymphoblastoma cell line 221 led to an increase in complex N-glycans on plasma membrane proteins. Biochemical and functional experiments revealed that increased glycosylation of ligands for NK activation receptors NKG2D and CD2 interfered with receptor binding and lysis of the target cells by NK cells. Analysis of ligands for NK receptors in *SPPL3*-KO cells revealed slower migration during gel electrophoresis, consistent with higher glycosylation. Pharmacological inhibition of the N-glycan pathway—at a step preceding branching mediated by MGAT transferases—reversed the slower gel migration of ligands and restored a normal sensitivity of *SPPL3*-KO cells to lysis by NK cells.

Notably, previous research has linked the loss of a chromatin regions encompassing *SPPL3* to reduced infiltration of T cells in lung adenocarcinoma [26]. This may be due to elevated glycosylation and disruptions in ligand-receptor interactions. Our study suggests that NK cells may also be impaired in this respect. Unlike mutations in specific ligands for NK receptors, the impact of *SPPL3* mutations in cancer is much broader. Therefore, we investigated the molecular basis for this association of SPPL3 and cancer. Using a sgRNA library focused on glycosylation for a secondary CRISPR screen in *SPPL3* knockout cells, we showed that enzymes promoting high branching of N-glycans and their elongation by addition of LacNAc were the main contributors to the evasion of *SPPL3*-deleted cells from lysis by NK cells.

NK cells lyse target cells mainly through two distinct pathways. One is the delivery of lytic effector molecules, such as granzymes, into target cells through membrane pores formed by polymerization of perforin. The second is through engagement of death receptor on target cells by TRAIL (TNFSF10), which is highly expressed on NK cells. Deletion of two death receptor genes *TNFRSF10A* and *TNFRSF10B* contributed to resistance in the primary genome-wide screen, suggesting that TNFSF10 on NK cells, which binds to both receptors, contributed to 221 cell death during selection in coculture. Over time, NK cells switch from rapid granzyme B-mediated killing to slower death-receptor-induced killing [27]. Given the long-term incubation and low effector to target ratio with primary NK cells used in the selection for resistant mutants, both pathways were given a chance to participate in elimination of target cells. A contribution of apoptosis to cell death of 221 cells in our screen was supported by deletions of *SRSF1* and *SRSF2* genes, which encode splicing factors of the Ser/Arg family (SRSF). SRSF1 promotes RNA splicing that prevents apoptosis, whereas SRSF2 promotes apoptosis [28, 29]. In the genome-wide screen *SRSF1* was among depleted sgRNAs, and *SRSF2* was among enriched sgRNAs, consistent with the anti- and pro-apoptosis function of SRSF1 and SRSF2, respectively.

Deletion of genes encoding ligands (CD58, MICB, PVR) of NK activation receptors also resulted in reduced sensitivity to NK cells. Their evasion score was lower because activation of NK cells requires co-engagement of several activation receptors [19]. The enhanced glycosylation of CD58 and MICB in *SPPL3*-KO cells interfered directly with binding of soluble receptors CD2 and NKG2D, respectively. Steric hindrance was likely responsible for this disruption considering that two distinct and specific ligand–receptor interactions were inhibited. Furthermore, receptor binding and activation of NK cell cytotoxicity were recovered after pharmacological inhibition of the N-glycosylation pathway at an early step, prior to the branching mediated by MGAT transferases. In addition, binding of the CD20 antibody rituximab was impaired on *SPPL3*-KO cells. This suggests that escape from treatment of mature B-cell non-Hodgkin’s lymphoma (NHL) and mature B-cell acute leukemia (B-AL) with rituximab can occur not only by downregulation of CD20 expression, but also by enhanced CD20 glycosylation. This should be an important consideration in the selection of antibodies for both monoclonal antibody therapies and single-chain variable fragments (scFvs) inserted into CARs.

Pathways previously known to protect or sensitize target cells facing an attack by NK cells were identified, providing a strong validation of our genetic screens. *B2M* knock-out on its own increased sensitivity to NK cells, and pathway analysis showed that several other genes involved in antigen processing and presentation pathway significantly scored in mediating resistance to NK cells. 221 cells lack expression of classical HLA class I molecules but do express HLA-E, a ligand for inhibitory receptor NKG2A. Likewise, the IFN-γ signaling pathway, which upregulates HLA class I expression and antigen processing was a significant protector against lysis by NK cells, as reported before [10, 30]. While such pathways protect cancer cells from recognition by NK cells, they have the opposite effect on recognition by CD8 T cells [14].

SPPL3 has been noted in other screens for sensitivity to cytotoxic T cells. Loss of *SPPL3* in the pre-B acute lymphoblastic leukemia (ALL) cell line NALM6 resulted in increased glycosylation of CD19 and a direct impairment of CAR T cell effector function and a reduction in their anti-tumor efficacy [31]. Another study reported that the response of CD8 T cells to HLA class I was impaired in *SPPL3*-KO cells through a different pathway, which involves an increase of glycosphingolipids in antigen-presenting cells. The accumulation of glycosyltransferases, such as B3GNT5, in *SPPL3*-KO cells increased the amount of glycosphingolipids at the plasma membrane, which sterically interfered with the accessibility of HLA class I to its receptors and diminished CD8^+^ T cell activation[32].

Having identified SPPL3 as a major regulator of sensitivity to NK cells, we went on to determine *how* a higher abundance of membrane-tethered intracellular glycosyltransferases impacts glycosylation of cell surface proteins and how this leads to target cell resistance. Was it mainly a quantitative effect due to higher numbers of glycosylated proteins and of glycosylation sites on proteins? Or was it more specific, due to changes in the type of N-glycans? To address this, *SPPL3*-KO cells were subjected to another round of selection by coculture with NK cells. This time a sgRNA library focused on genes that control glycosylation pathways was used to identify the type of glycans that may be responsible for resistance to NK cells. Two independent screens with this library identified MGAT2, which adds GlcNAc on the second branch after the trimming of high mannose precursors, and B3GNT2, a transferase specialized in the extension of branched N-glycans by adding GlcNAc in a β1,3 linkage to an unsialylated terminal galactose [24]. B3GNT2 has also been identified as a driver of resistance of human melanoma cells to T-cell mediated cytotoxicity [33]. B3GNT2 prefers to elongate the fourth branch, which is added by MGAT5 [34, 35]. B3GNT2 competes with sialyltransferases for the same substrate, namely one of the 4 terminal Gal on tetra-antennary N-glycans. Sialyltransferases get a head start, as they sialylate any terminal Gal, including those on partially branched N-glycans, while B3GNT2 prefers tetra-antennary N-glycans. Remarkably, the resistance to NK-mediated lysis conferred by *SPPL3*-KO was further *increased* by deletion of *ST6GAL1*. As ST6GAL1 specializes in α2,6-linked sialylation of terminal Gal on branched N-glycans [36], its absence provides more opportunities for branch extension by B3GNT2.

In addition to *MGAT2*, the other three *MGAT* genes 1, 4, and 5 required for synthesis of tetra-antennary N-glycans were also detected in our screen, providing further evidence that high N-glycan branching contributes to the resistant phenotype. Expression of *MGAT5* has been linked to cancer [34, 35, 37] and inhibits killing of pancreatic adenocarcinoma by CAR-T cells targeting CD44 [38]. In contrast to *MGAT2*, knockout of *MGAT3* conferred additional *resistance* against NK-mediated killing to *SPPL3* knockout cells. MGAT3 transfers a GlcNAc in a β1,4 linkage to the core β-mannose of N-glycan, a step which is referred to as a bisecting GlcNAc. Addition of a bisecting GlcNAc inhibits further branching by MGAT4 and MGAT5 [34, 39]. Therefore, the absence of MGAT3 promotes MGAT4 and MGAT5-mediated addition of the third and fourth N-glycan antenna, thereby increasing the number of substrates for B3GNT2-mediated elongation and the resistance of target cells to NK-mediated lysis.

N-glycan species with LacNAc extensions identified by our glycomic analysis included four that carried one LacNAc, and one each with extensions of 2, 4, and 6 LacNAc units, all of which were enriched in *SPPL3*-KO cells. Our glycoproteomic analysis demonstrated the presence of tetra-antennary N-glycans that included at least one LacNAc extension. They were identified on abundant proteins only, which included CD48 and ICAM-1. These extended N-glycans were significantly enriched in *SPPL3*-KO cells. CD48 is the ligand of NK receptor CD226 (2B4, SLAMF4). High branching and elongation of CD48 N-glycans could be relevant to NK cytotoxicity toward adult T cell leukemia and lymphoma (ATLL), which depends on CD48 expression [40].

In this study, we demonstrate that SPPL3 functions as a primary determinant of interactions between NK cells and tumor cells, with profound implications for cancer biology. We identified the specific steps in the N-glycosylation pathway in target cells that contributed the most to resistance against NK cells, namely high branching of N-glycans that are further elongated by addition of LacNAc. Malignant B-cells and other tumor cells that upregulate transferases in this pathway may acquire resistance against cytotoxic T and NK cells. Downregulation or mutations of the *SPPL3* gene can promote tumor cell escape due to intracellular accumulation of several glycosyltransferases [26]. Our study supports the feasibility of incorporating glycosylation pathway inhibitors into cancer treatment regimens to further enhance the efficacy of cancer therapy. Moreover, CAR-T and CAR-NK cells are at the forefront of cancer immunotherapy and their effectiveness relies on specific recognition of tumor cell antigens. In this regard, a major concern is the diminished binding of tumor-specific antibodies in the context of monoclonal antibody therapies and single-chain variable fragments (scFv) that are built into CARs. It is therefore critical to identify antibody epitopes that are not shielded by large glycans on tumor antigens to ensure the targeted destruction of tumor cells by monoclonal antibody and CAR therapies.

## Materials and Methods

### Key reagents

| Reagent type | Designation | Source | Identifier | Clone name |
| --- | --- | --- | --- | --- |
| Antibody | Anti-human CD107a FITC | BD Biosciences | 555800 | H4A3 |
| Antibody | Anti-human CD56 BV421 | BD Biosciences | 562751 | NCAM16.2 |
| Antibody | Anti-human SPPL3 | Millipore Sigma | MABS1910 | 7F9 |
| Antibody | Anti-human FBXO11 polyclonal | NOVUS | NB100-59826 |  |
| Antibody | Anti-human CD58 | BD Biosciences | 555919 | 1C3 |
| Antibody | Anti-human CD58 PE | BD Biosciences | BDB555921 | 1C3 |
| Antibody | Anti-human MICB PE | R&D Systems | FAB1599P | 236511 |
| Antibody | Anti-human MICB polyclonal | R&D Systems | AF1599 |  |
| Antibody | Anti-human TRAIL-R1 PE | BD Biosciences | 564180 | S35-934 |
| Antibody | Anti-human TRAIL-R2 PE | BD Biosciences | 565499 | YM366 |
| Antibody | Anti-human CD48 PE | BD Biosciences | 552855 | TU145 |
| Antibody | Anti-human ICAM1 | BD Biosciences | 555511 | HA58 |
| Antibody | Anti-human HLA-E | BioLegend | 342604 | 3D12 |
| Antibody | Anti-human MGAT5 | R&D Systems | MAB5469 | 706824 |
| Antibody | Anti-human actin | Santa Cruz | sc-8432 | C-2 |
| Antibody | Anti-HA high affinity | Millipore Sigma | 11867423001 | 3F10 |
| Recombinant protein | Human CD2 Fc Chimera Protein | R&D Systems | 1856-CD |  |
| Purified protein | Human NKG2D Fc Chimera Protein | R&D Systems | 1299-NK-050 |  |
| Lectin | Concanavalin A (Con A), FITC | Vector labs | FL-1001-25 |  |
| Lectin | Sambucus Nigra Lectin (SNA), FITC | Vector labs | FL-1301-2 |  |
| Lectin | Phaseolus Vulgaris Erythroagglutinin (PHA-E), FITC | Vector labs | FL-1121-2 |  |
| Lectin | Lycopersicon esculentum Lectin (LEL), FITC | Vector labs | FL-1171-1 |  |
| Chemical inhibitor | Kifunensine | Millipore Sigma | K1140 |  |
| Chemical inhibitor | Swainsonine | Millipore Sigma | S8195 |  |
| CRISPR library | GeCKO V2 human CRISPR knockout | Addgene | 1000000049 |  |
| CRISPR library | Human GlycoGene CRISPR pooled library | Addgene | 140961 |  |
| Fluorescent Dye | PKH26 Red Fluorescent Cell Linker Kit | SIGMA | PKH26GL |  |
| Fluorescent Dye | PKH67 Green Fluorescent Cell Linker Kit | SIGMA | PKH67GL |  |
| Counting beads | 123count eBeads™ | ThermoFisher | 01-1234-42 |  |
| Commercial kit | Dead Cell Removal Kit | Miltenyi Biotec | 130-090-101 |  |
| Commercial kit | In-Fusion HD cloning kit | Clontech Laboratories | 639648 |  |
| Commercial kit | EasySep Human NK Cell Enrichment Kit | STEMCELL Technologies | 19055 |  |
| Commercial kit | EasySep™ Human CD56 Positive Selection Kit | STEMCELL Technologies | 17855 |  |
| PCR polymerase | NEBNext® High-Fidelity 2X PCR Master Mix | New England Biolabs | M0541L |  |

### Cells

Peripheral blood samples from healthy US adults were obtained from the NIH Department of Transfusion Medicine under an NIH Institutional Review Board-approved protocol (99-CC-0168) with informed consent. NK cells were isolated using negative selection (Stemcell Technologies). The NK cells were then suspended in IMDM (Gibco) containing 10% human serum (Valley Biomedical) and were used within 4 days. To generate IL-2-activated NK cells, the freshly isolated NK cells were cultured together with irradiated autologous feeder cells in the presence of 10% purified IL-2 (Hemagen), 100 units/mL recombinant IL-2 (Roche), and 5 μg/mL PHA (Sigma), all in OpTimizer (Invitrogen) medium. Subsequently, expansion was carried out in the same medium, but without the PHA and feeder cells. 221 cells were cultured in RPMI 1640 medium (Gibco) with the addition of 10% heat-inactivated fetal calf serum (Gibco) and 2 mM L-Glutamine (Gibco). To block the glycosylation pathway, cells were either treated with 4 μg/mL kifunensine or 10 μg/mL swainsonine for 24 hours before lectin staining or functional assays.

### Plasmids and lentivirus production

The sgRNAs that targeted specific genes were synthesized by Integrated DNA Technologies (IDT) and cloned into the BsmBI restriction sites of the LentiGuide-Puro vector (Addgene #52963) using the In-Fusion HD cloning kit (Clontech). To make SPPL3 and B3GNT2 double knockout cells, sgSPPL3-1 was cloned into a LentiGuide-hygro vector, which was modified from the LentiGudie-Puro vector by replace the puromycin-resistant gene with a hygromycin-resistant gene using the In-Fusion cloning kit. The following sgRNA sequences were used in this study, sgSPPL3-1 (CTTGTTAAATACTGGCACAT), sgSPPL3-2 (CGAGTAGGTCTGCTCCGCCA), FBXO11 (ACTTCAACTACAGAAAACTT), sgNT (TCCTGCCAAGAAACACCCTT), sgMICB (CACCTGCAGCGAGGTCTCAG), sgCD58 (CATGTTGTAATTACTGCTAA), sgTNFRSF10B (ACACATTCGATGTCACTCCA), sgTNFRSF10A (ACACACTCGATGTCACTCCA) and sgB3GNT2 (GTTCCTCTACTCCGGCCACC). To prepare for lentivirus production, low-passage Lenti-X 293T cells (Clontech) were seeded into a T75 flask and transfected the following day using PEI Max (Polyethylenimine). The transfection procedure involved mixing 1.2 μg of pMD2.G plasmid, 2.3 μg of psPAX2 plasmid, 4.6 μg of LentiGuide-Puro plasmid, and 217 μl of serum-free DMEM in a Falcon tube. Then, 65 μl of PEI Max 40K (Polysciences) stock solution (1 mg/ml) was added and briefly vortexed. After 10 minutes, 8.6 ml of DMEM media containing 10% FCS was added to the tube. The culture medium for the 293T cells was replaced with the fresh medium containing the transfection mixture. Supernatants were collected two days after transfection, passed through a 0.45 μM filter, and stored at −80°C. The virus was either used directly or concentrated using PEG-it (System Biosciences) before use. To transduce human cell lines, the lentivirus was combined with polybrene to a final concentration of 8 μg/ml and added to the cells, which were then incubated for 2 days. The cells were subsequently centrifuged at 1200 rpm for 10 minutes and resuspended in complete medium with puromycin at pre-titrated concentrations.

### Genome-wide CRISPR screen

The 221 cells were transduced with LentiBlast-Cas9 and selected by the addition of 10 μg/ml blasticidin. The GeCKO V2 human CRISPR knockout library from Addgene was then introduced into Endura Electrocompetent Cells by means of electroporation using a Bio-Rad Gene Pulser. The expanded CRISPR plasmid libraries were purified using Maxi-Prep and used to produce lentivirus. The lentivirus titer was determined using a previously described method [41]. The 221 cells that expressed Cas9 were transduced with the GeCKO V2 lentivirus library at a low multiplicity of infection (MOI) of 0.3 and selected with puromycin for 10 days. At least 180 x 10^6^ cells were used for transduction to ensure a coverage of >500 cells per sgRNA. The transduced 221 cells were selected and culture with 1 μg/ml puromycin for 10 days then incubated with IL-2-activated NK cells at a ratio of 1:5 and incubated for 48 to 72 hours. The percentage of surviving 221 cells was monitored and, if necessary, additional NK cells were added until 10% of the 221 cells remained. To recover the surviving 221 cells, the dead cells were first removed with a Dead Cell Removal Kit and then the NK cells were depleted using the EasySep Human CD56 Positive Selection Kit (Stemcell Technologies). The control 221 cells were kept in the same culture conditions but were not exposed to NK cells. The screen was performed with two biological repeats. The genomic DNA extraction and gRNA cassette amplification were carried out using a previously described method [41]. The amplified libraries were then multiplexed and analyzed on a NextSeq 500 with 75-bp single-end reads. The analysis of gRNA enrichment/depletion was performed using MAGeCK [42] MLE. This pipeline calculated the individual sgRNA read counts in libraries from both control and surviving 221 cells and then normalized the read counts of individual gRNAs based on control sgRNAs included in the library. The read counts of enriched gRNAs increased in surviving 221 cells compared to control 221 cells, while the read counts of depleted sgRNAs decreased in surviving 221 cells. The score of each gene represents the normalized log-fold change of all gRNAs targeting that gene.

### Glycosylation-focused secondary CRISPR screen

Production of lentivirus from glycosylation-focused CRISPR library (glycoGene)[23] was described above. This glycoGene library consisted of 3637 sgRNAs targeting 347 genes involved in the glycosylation pathways. The library backbone constitutively expresses BFP for selection. SPPL3-deleted or sgNT 221 cells expressing LentiBlast-Cas9 were transduced with library virus at a low MOI of 0.3 for 2 days and cultured for an additional 2 days. At least 2 x 10^6^ BFP positive 221 cells were sorted for each biological repeat to ensure coverage of >500 cells per sgRNA. Sorted cells were treated with doxycycline (dox) for 10 days to induce sgRNAs expression and genomic editing. Then, cells were co-incubated with NK cells at low E:T ratios and selected until 10% to 20% of the 221 cells remained alive. Dead cells were removed with a Dead Cell Removal Kit (Miltenyi biotec). At least 4 x 10^6^ million surviving cells were collected for DNA extractions and library amplification. Genomics DNA extractions were conducted using the PureLink Genomic DNA Mini Kit (Invitrogen). Library amplification was carried out by a two-step approach as described elsewhere [23]. Barcoded libraries were pooled and sequenced on an illumina MiSeq platform with 150 bp paired end reads.

### Cytotoxicity and degranulation assay

The cytotoxicity assay was carried out using a flow cytometry-based approach. In brief, 221 cells were labeled with either PKH67-green or PKH26-red membrane dyes, then rinsed with complete growth medium. The IL-2-expanded NK cells were then combined with the pre-labeled 221 cells and incubated at 37°C in IMDM media supplemented with 10% FBS for overnight. The number of viable 221 cells was determined by counting beads using a flow cytometer. For ADCC assays, 221 cells were pre-incubated with 10 μg/mL rituximab for 15 min at room temperature before adding NK cells for lysis. Assessment of degranulation was performed using resting human NK cells. The target cells were combined with NK cells and incubated for 2 hours at 37°C. The cells were then stained with Live/Dead-NIR (Thermo Fisher), anti-CD56-Bv421 (BD 562751), and anti-CD107a-FITC (BD 555800). Subsequently, flow cytometry was used to analyze the samples.

### Flow cytometry

For immune staining before flow cytometry, cells were incubated with premixed fluorophore-conjugated antibodies diluted in FCAS buffer (PBS, 0.5 % BSA) at 4°C for 30 min. Cells were washed after staining and analyzed on an LSR II (BD Biosciences) or LSRFortessa X-20 (BD Biosciences). Data were analyzed with FlowJo (FlowJo, LLC). For lectin staining, cells were incubated with fluorophore-conjugated lectins diluted in HBSS buffer (ThermoFisher Scientific) for 30 min at 4°C. Cells were washed and analyzed on a flow cytometer.

### Western blot

To prepare samples for western blot, the cell pellets were resuspended in RIPA buffer containing 50 mM Tris-HCl (pH 8.0), 150 mM NaCl, 1.0% NP-40, 0.5% SDS, 0.5% sodium deoxycholate, cOmplete protease inhibitor cocktail (Roche), and PhosSTOP phosphatase inhibitor cocktail (Millipore Sigma). The samples were then lysed on ice for 30 minutes and the cellular debris was removed by spinning at 12,000 rpm for 10 minutes. The protein lysates were mixed with 4X NuPAGE LDS-PAGE sample buffer (Invitrogen), separated on 4-12% Bis-Tris gels, transferred to PVDF membranes, and probed with the relevant antibodies. The chemiluminescent signal was detected using a ChemiDoc Imaging system (Bio-Rad).

To detect endogenous SPPL3 by western blot, a membrane extraction procedure was carried out [43]. Briefly, cells were resuspended in hypotonic buffer kept at 4°C and sheared using a needle. The nuclei were sedimented by centrifugation at 2400g for 5 minutes at 4°C, and the resulting membrane pellets were obtained by centrifugation at 21,000g or 100,000g. The membrane pellets were then washed twice with carbonate buffer (0.1 M Na2CO3, 1 mM EDTA, pH 11.3), followed by washing with STE buffer (150 mM NaCl, 50 mM Tris, 2 mM EDTA, pH 7.6). Finally, membrane pellets were lysed in RIPA buffer and analyzed by western blot.

### RNA-seq and data analysis

Cells were harvested in Trizol and combined with 200μl of 1-Bromo-3-chloropropane (Millipore Sigma), mixed vigorously, and centrifuged at 16,000 x g for 15 min at 4°C. RNA containing aqueous phase (600 μl) was collected and passed through QiaShredder column (Qiagen, Valencia, CA) at 21,000 x g for 2 minutes to homogenize any remaining genomic DNA in the aqueous phase. Aqueous phase was combined with 600uL of RLT lysis buffer (Qiagen, Valencia, CA) with 1% beta mercaptoethanol (MilliporeSigma, St. Louis, MO) and RNA was extracted using Qiagen AllPrep DNA/RNA 96 kit (Valencia, CA). An additional on-column DNase I treatment was performed during RNA extraction. All sample processing was performed using amplicon-free reagents and tools in aerosol-resistant vials. RNA quality was analyzed using Agilent 2100 Bioanalyzer (Agilent Technologies, Santa Clara, CA). RNA samples were quantitated individually using a Qubit RNA High Sensitivity assay (Thermofisher, Waltham, MA) to create 1 μg aliquots for each. Samples were then processed using Truseq Stranded mRNA Library preparation kit (Illumina Inc., San Diego, CA) and single-end indexed for multiplexing. The resulting libraries were titrated using Kapa Library Quantification kit (Universal) (Roche, Basel, Switzerland) and measured on a Touch96 RTPCR instrument (Bio-Rad Laboratories, Hercules, CA). The samples were diluted to 2 nM working stocks and pooled together for sequencing on Illumina NextSeq MID-Output runs for 75 cycles in both directions and an additional 6 cycles to read the index. RNA-seq reads were compiled and filtered to remove any reads with PHRED scores less than 10 and aligned to the human hg38 genome using bowtie2. FeatureCounts was used to determine reads for annotated genes, and differential expression analysis was conducted using edgeR.

### N-glycoproteomic sample preparation, MS analysis and database search

The site-specific N-glycoproteomic experiment was performed as previously described [44]. Briefly, cell pellets were lysed in the lysis buffer containing 4% SDS (w/v), 50 mM HEPES, pH 8.0 and sonicated for 20 min (30 s on, 30 s off) using Bioruptor at 4 °C. After centrifugation at 14,000 × g for 15 min, the supernatants were collected, and the protein amounts were measured using Pierce BCA Protein Assay Kit (Thermo Scientific). 400 μg protein from each sample was reduced and alkylated with 10 mM TCEP (0.5 M stock, Thermo Scientific) and 20 mM IAM at 37 °C for 60 min in the dark. A mixture of Sera-Mag SpeedBeads (GE Healthcare, cat.no. 45152101010250; cat.no. 65152105050250) was rinsed twice with water and then added to protein lysates at the working ratio of 10:1 (wt/wt, beads to proteins). After adding acetonitrile (ACN) to a final percentage of 70% (v/v), the beads and proteins were incubated for 10 min at room temperature off the rack, followed by resting on magnetic rack for 2 min to remove the supernatant. The beads were washed three times with 90% (v/v) ACN, resuspended in 50 mM HEPES (pH 8.0) containing sequencing grade modified trypsin (1:40 of enzyme-to-protein ratio), and incubated at 37°C overnight in a ThermoMixer with mixing at 1000 rpm. The resulting digested peptides in the supernatant were collected, followed by labelling with TMTsixplex™ isobaric label Reagents according to the manufacturer’s instruction (Thermo Scientific).

The dried TMT-labelled peptides were redissolved in loading buffer (80% (v/v) ACN and 1% TFA) and loaded five times to spin-columns self-packed with the Ultimate Hydrophilic Interaction Liquid Chromatography (HILIC) Amphion II beads (5 μm, Welch, Lot. no. 7301.26). After three washes with washing buffer (75% (v/v) ACN, 1% TFA), the retained TMT-labeled glycopeptides were eluted with 100 μL 0.1% TFA twice and dried in a SpeedVac concentrator. Subsequently, basic reverse phase fractionation for enriched glycopeptides was performed on an Agilent 1100 series high-performance liquid chromatography (HPLC) system installed with a XBridge C18 column (3.5 μm particles, 1.0 mm × 150 mm, Waters). Peptides were collected in a time-based mode from 6 to 64 min and pooled into 6 concatenated fractions.

LC-MS analysis was performed on an Orbitrap Fusion Lumos Tribrid Mass Spectrometer (Thermo Scientific) coupled to a Dionex UltiMate 3000 UHPLC system (Thermo Scientific). The MS instrument settings were described briefly in the following. MS1 settings: Orbitrap Resolution-120 k, Mass Range (m/z)-350-2000, Maximum IT-50 ms, AGC target-5e5, RF Lens-60%, Precursor selection range (m/z)-700-2000; MS2 settings: Isolation window-2 m/z, Scan range mode-define first mass-132, Activation type-HCD, Collision energy-25, Detector type-Orbitrap, Orbitrap resolution-15 K, Maximum IT-150 ms, AGC target-5e5; MS3 settings: Precursor selection range-700-2000, Number of Notches-10, First mass (m/z)-120, Activation type-HCD, Collision energy (%)-35, Detector type-Orbitrap, Orbitrap resolution-15 K, Maximum IT-350 ms, AGC target-5e5, Number of Dependent Scans-10.

All raw files were processed by GlycoBinder [44]. Main parameters used for GlycoBinder include fully specific trypsin digestion with maximal two missed cleavage and mass tolerance for precursors and fragment ions of 10 and 20 ppm, respectively. Cysteine carbamidomethylation and TMT6 on peptide N-termini and lysine residues were set as fixed modifications and methionine oxidation was set as a variable modification. The reviewed human protein database was downloaded from Swiss-Prot (October 2020, human, 20,370 entries). GlycoBinder propagates all glycopeptide-to spectra matches and directly reported the quantification of glycosylation sites, glycan compositions, and glycoforms in separate txt files.

All identified N-glycopeptides and their glycan compositions were classified into 6 glycan types based on the number of different monosaccharide moieties, including hexose (Hex), N-acetylhexosamine (HexNAc), N-acetylneuraminic acid (Neu5Ac), and fucose (Fuc). “Complex” type corresponds to the glycan composition of HexNAc (≥4)-Hex(≥3)-NeuAc(any)-Fuc(any). “A4/A3B” represents 4 potential number of branches or 3 branches with a bisect, corresponding to the glycan composition of HexNAc (≥6)-Hex(≥3)-NeuAc(any)-Fuc(any).

## Acknowledgements

This research was supported by the Division of Intramural Research, National Institute of Allergy and Infectious Diseases (NIAID), NIH. Y.J. and K.-T.P. were supported in part by the LOEWE Center Frankfurt Cancer Institute (FCI) funded by the Hessen State Ministry for Higher Education, Research and the Arts [III L 5-519/03/03.001-(0015)]. S.S. was supported by a grant from the German Research Foundation (Deutsche Forschungsgemeinschaft [DFG] SCHE 2120/1-1). We thank the Sequencing Facility of the National Cancer Institute for next-generation sequencing of the genome-wide CRIPSR screen library, C. Martens and his team at the Genomics Research Section, Genomics Research at the Research Technologies Branch (RTB), NIAID for their input and the NGS sequencing of the secondary CRIPSR screen libraries and RNA-seq, P. Azadi and B. Kumar at the Complex Carbohydrate Research Center, University of Georgia, for the glycomics analysis, and S. Rajagopalan for discussions and comments on the manuscript.

## Author contributions

X.Z. and E.O.L. conceived the project. X.Z., J.W. and Y.J. performed the experiments. X.Z., M.V. and K.-T.P. provided the methodology for performing the experiments. X.Z, Y.J. and K.-T.P. contributed formal analysis. X.Z. and E.O.L. wrote the manuscript. X.Z., E.O.L., M.V., F.M., J.W., Y.J. and S.S. edited the manuscript. E.O.L and K.-T.P. supervised the study. E.O.L, H.U. and K.-T.P. acquired funding.

## Data Availability

CRISPR screen and RNA-seq data in this paper are available at GEO under the accession number GSE228188. The mass spectrometry-based glycoproteomics data has been deposited to the ProteomeXchange Consortium via the PRIDE [45] partner repository with the dataset identifier PXD040769 with Username, Password: 0FUpzxfj during the peer review process.

---

**Fig. S1.**
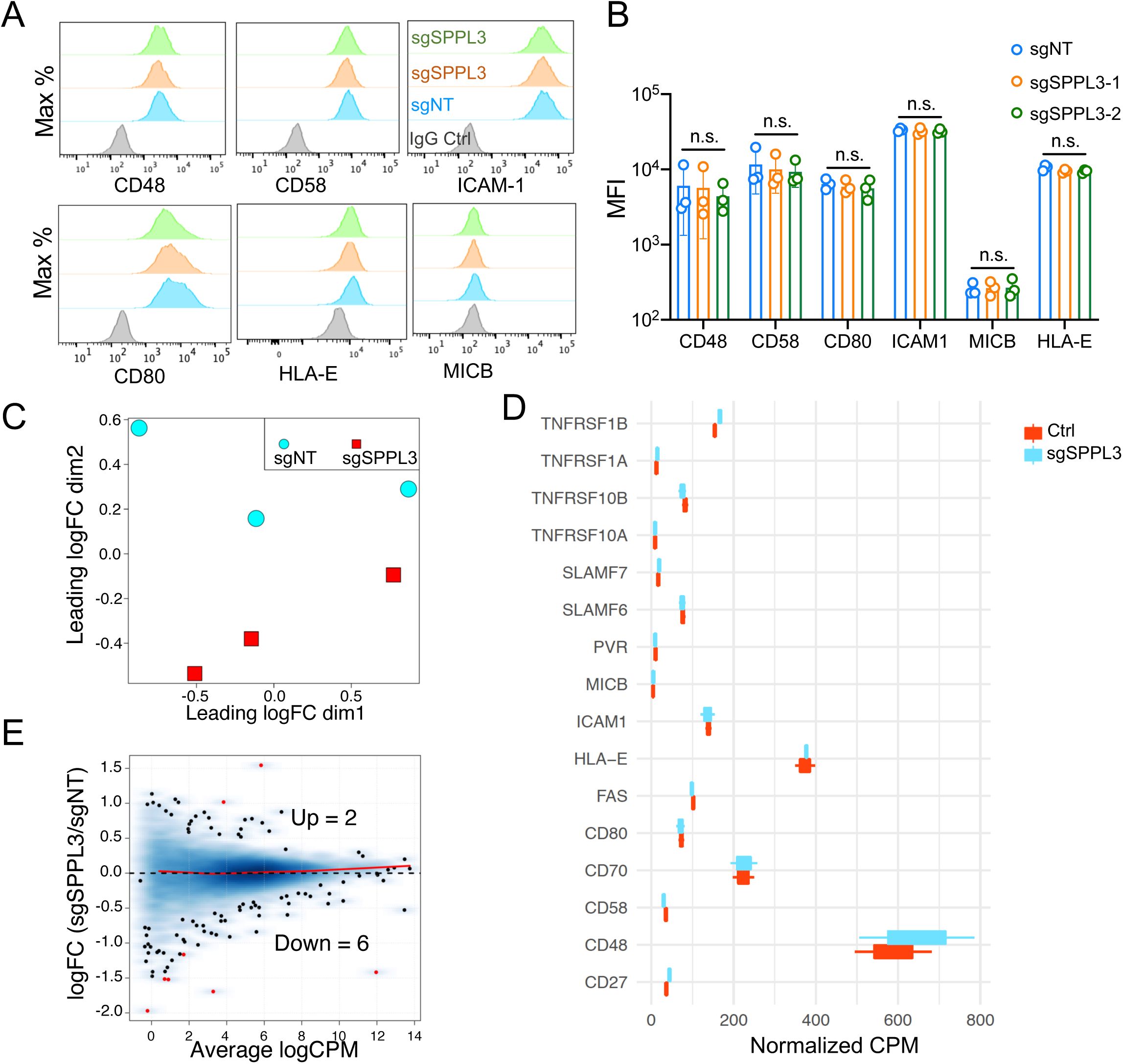
Impact of SPPL3 deletion on surface expression of ligands for NK receptors and their transcription. (A) Representative histograms showing surface staining of indicated ligands for NK receptors on 221 cells expressing sgNT or sgSPPL3. (B) Statistics of mean fluorescence intensity (MFI) from 3 experiments carried out as in (A). Data shown as mean ± SEM (n=3, one-way ANOVA test, n.s. not significant). (C) PCA plot shows sample clustering of RNA-seq data from sgNT and sgSPPL3 221 cells. (D) Transcription level of the indicated genes encoding ligands of NK receptors in sgNT and sgSPPL3 221 cells. (E) Mean-difference (MD) plot of global transcriptomic changes in sgSPPL3 221 cells compared to sgNT cells.

**Fig S2.**
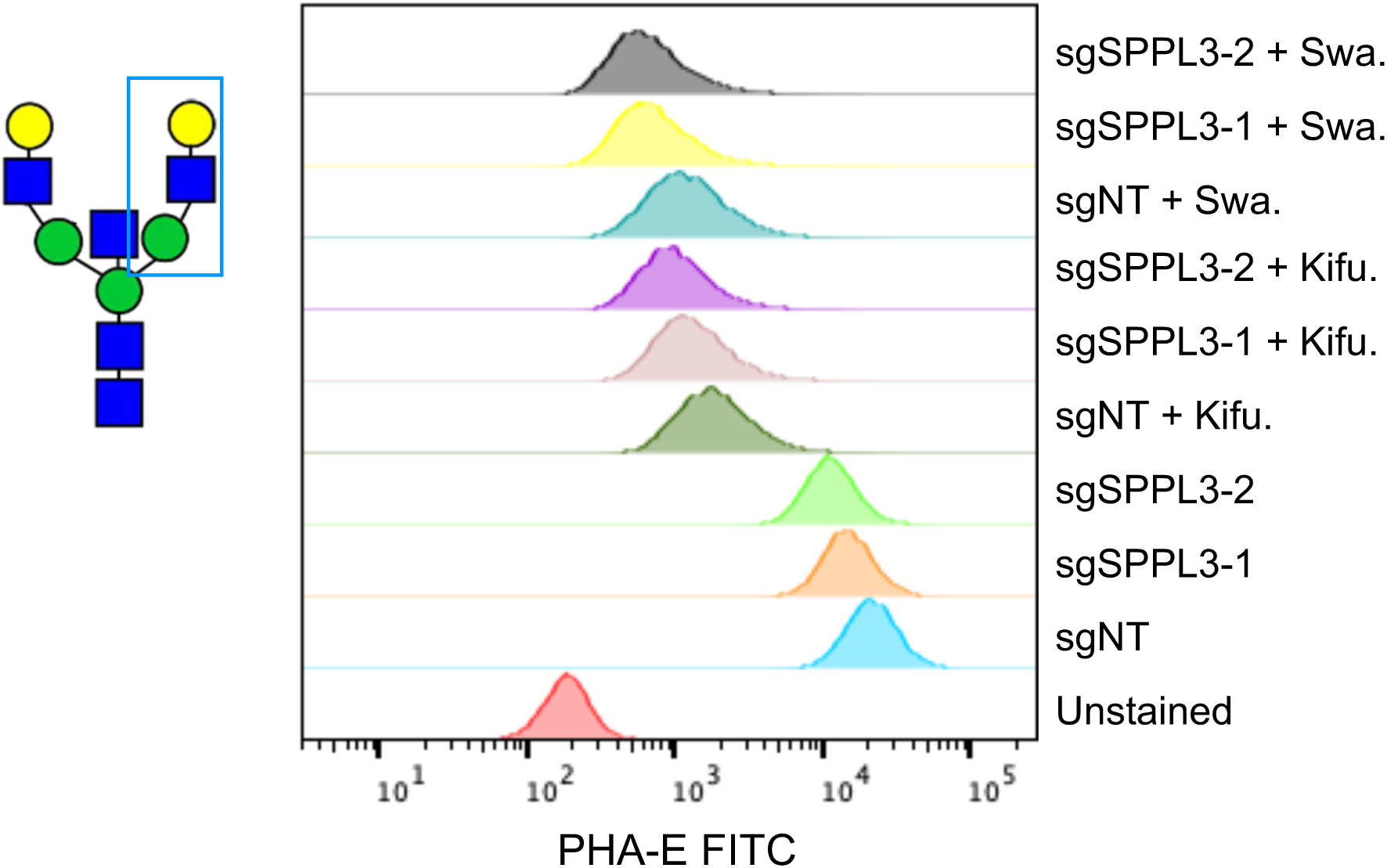
Representative histograms of PHA-E staining of 221 cells expressing sgNT or sgRNAs targeting SPPL3. Cells were non-treated, treated with kifunensine (Kifu.) or swainsonine (Swa.).

**Fig. S3.**
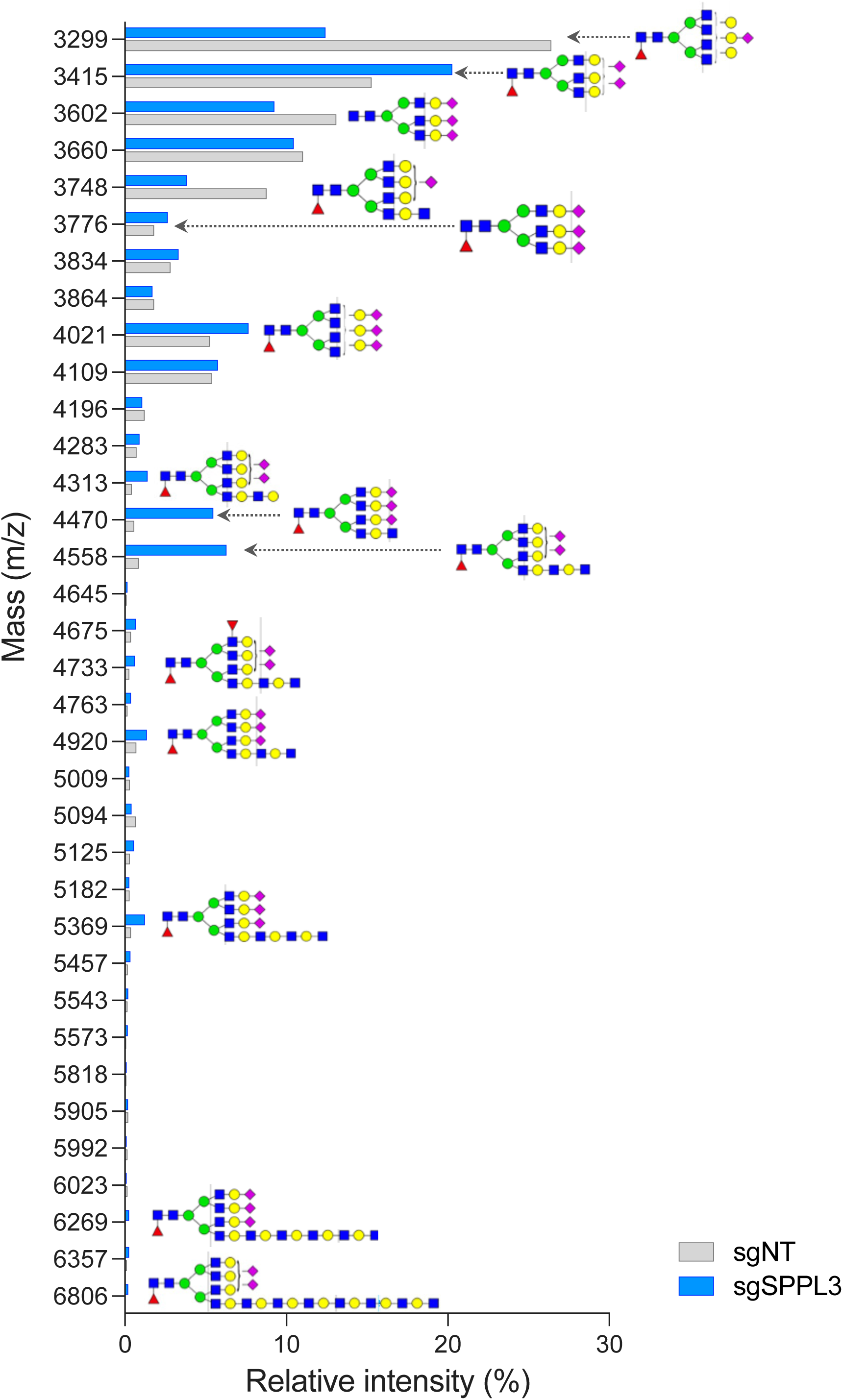
Glycomic analysis of high mass N-glycans on 221 cells expressing sgNT (grey) or sgSPPL3 (blue). Displayed are high mass N-glycans (>3298) detected in sgSPPL3-221 cells and compared with their abundance in sgNT-221 cells. All of them are at least tri-antennary. The two glycan forms that were reduced the most in sgSPPL3-221 cells are tetra-antennary forms with 2 unmodified Gal, neither sialylated nor extended with GlcNAc (m/z 3299 and 3748). The most enriched form was tetra-antennary with 3 sialylated Gal and 1 GlcNAc-elongated Gal (m/z 4470). Addition of Gal onto GlcNAc by a B4GALT generates LacNAc. The first tri-sialylated N-glycan with a LacNAc extension is at m/z 4920. This form has already acquired a GlcNAc on top of the first LacNAc. Further LacNAc elongations were detected as enriched N-glycans carrying 2 LacNAc (m/z 5369) and a low abundance 4 LacNAc (m/z 6269). The second most enriched (m/z 4558) has only 2 sialylated Gal, a LacNAc elongation, and a terminal GlcNAc. This terminal GlcNAc could be either on a separate branch or at the tip of the elongated branch, as depicted. This N-glycan could the precursor of the very large but low abundance N-glycan elongated with 6 LacNAc (m/z 6806). Note that the 6 LacNAc extensions could be distributed among the two unsialylated Gal. Note that the first glycan form carrying a LacNAc extension is at m/z 4313, which has two sialylated Gal, one free Gal and one LacNAc. It is a precursor of m/z 4558. It is formally possible that the terminal GlcNAc found on several N-glycan forms was added at an earlier step to the first mannose by MGAT3 (referred to as a bisection). It is however unlikely because bisection inhibits further branching by MGAT4 and MGAT5 [39].

## References

1. Long, E.O., et al., Controlling natural killer cell responses: integration of signals for activation and inhibition. Annu Rev Immunol, 2013. 31: p. 227–58.

2. Myers, J.A. and J.S. Miller, Exploring the NK cell platform for cancer immunotherapy. Nat Rev Clin Oncol, 2021. 18(2): p. 85–100.

3. Morvan, M.G. L.L. Lanier, NK cells and cancer: you can teach innate cells new tricks. Nat Rev Cancer, 2016. 16(1): p. 7–19.

4. Chiossone, L., et al., Natural killer cells and other innate lymphoid cells in cancer. Nat Rev Immunol, 2018. 18(11): p. 671–688.

5. Barry, K.C., et al., A natural killer-dendritic cell axis defines checkpoint therapy-responsive tumor microenvironments. Nat Med, 2018. 24(8): p. 1178–1191.

6. Bottcher, J.P., et al., NK Cells Stimulate Recruitment of cDC1 into the Tumor Microenvironment Promoting Cancer Immune Control. Cell, 2018. 172(5): p. 1022–1037 e14.

7. Souza-Fonseca-Guimaraes, F., J. Cursons, and N.D. Huntington, The Emergence of Natural Killer Cells as a Major Target in Cancer Immunotherapy. Trends Immunol, 2019. 40(2): p. 142–158.

8. Liu, E., et al., Use of CAR-Transduced Natural Killer Cells in CD19-Positive Lymphoid Tumors. N Engl J Med, 2020. 382(6): p. 545–553.

9. Huntington, N.D., J. Cursons, and J. Rautela, The cancer-natural killer cell immunity cycle. Nat Rev Cancer, 2020. 20(8): p. 437–454.

10. Zhuang, X., D.P. Veltri, and E.O. Long, Genome-Wide CRISPR Screen Reveals Cancer Cell Resistance to NK Cells Induced by NK-Derived IFN-gamma. Front Immunol, 2019. 10: p. 2879.

11. Dubrot, J., et al., In vivo CRISPR screens reveal the landscape of immune evasion pathways across cancer. Nat Immunol, 2022. 23(10): p. 1495–1506.

12. Freeman, A.J., et al., Natural Killer Cells Suppress T Cell-Associated Tumor Immune Evasion. Cell Rep, 2019. 28(11): p. 2784–2794 e5.

13. Sheffer, M., et al., Genome-scale screens identify factors regulating tumor cell responses to natural killer cells. Nat Genet, 2021. 53(8): p. 1196–1206.

14. Patel, S.J., et al., Identification of essential genes for cancer immunotherapy. Nature, 2017. 548(7669): p. 537–542.

15. Challa-Malladi, M., et al., Combined genetic inactivation of beta2-Microglobulin and CD58 reveals frequent escape from immune recognition in diffuse large B cell lymphoma. Cancer Cell, 2011. 20(6): p. 728–40.

16. Klanova, M., et al., Prognostic Impact of Natural Killer Cell Count in Follicular Lymphoma and Diffuse Large B-cell Lymphoma Patients Treated with Immunochemotherapy. Clin Cancer Res, 2019. 25(15): p. 4634–4643.

17. Shimizu, Y., et al., Transfer and expression of three cloned human non-HLA-A,B,C class I major histocompatibility complex genes in mutant lymphoblastoid cells. Proc Natl Acad Sci U S A, 1988. 85(1): p. 227–31.

18. Kuhn, P.H., et al., Secretome analysis identifies novel signal Peptide peptidase-like 3 (Sppl3) substrates and reveals a role of Sppl3 in multiple Golgi glycosylation pathways. Mol Cell Proteomics, 2015. 14(6): p. 1584–98.

19. Bryceson, Y.T., et al., Synergy among receptors on resting NK cells for the activation of natural cytotoxicity and cytokine secretion. Blood, 2006. 107(1): p. 159–66.

20. Bryceson, Y.T., H.G. Ljunggren, and E.O. Long, Minimal requirement for induction of natural cytotoxicity and intersection of activation signals by inhibitory receptors. Blood, 2009. 114(13): p. 2657–66.

21. Voss, M., et al., Shedding of glycan-modifying enzymes by signal peptide peptidase-like 3 (SPPL3) regulates cellular N-glycosylation. EMBO J, 2014. 33(24): p. 2890–905.

22. Batlevi, C.L., et al., Novel immunotherapies in lymphoid malignancies. Nat Rev Clin Oncol, 2016. 13(1): p. 25–40.

23. Zhu, Y., et al., A GlycoGene CRISPR-Cas9 lentiviral library to study lectin binding and human glycan biosynthesis pathways. Glycobiology, 2021. 31(3): p. 173–180.

24. Kadirvelraj, R., et al., Comparison of human poly-N-acetyl-lactosamine synthase structure with GT-A fold glycosyltransferases supports a modular assembly of catalytic subsites. J Biol Chem, 2021. 296: p. 100110.

25. Truberg, J., et al., Endogenous tagging reveals a mid-Golgi localization of the glycosyltransferase-cleaving intramembrane protease SPPL3. Biochim Biophys Acta Mol Cell Res, 2022. 1869(11): p. 119345.

26. Bentham, R., et al., Using DNA sequencing data to quantify T cell fraction and therapy response. Nature, 2021. 597(7877): p. 555–560.

27. Prager, I., et al., NK cells switch from granzyme B to death receptor-mediated cytotoxicity during serial killing. J Exp Med, 2019. 216(9): p. 2113–2127.

28. Anczukow, O., et al., The splicing factor SRSF1 regulates apoptosis and proliferation to promote mammary epithelial cell transformation. Nat Struct Mol Biol, 2012. 19(2): p. 220–8.

29. Kedzierska, H. and A. Piekielko-Witkowska, Splicing factors of SR and hnRNP families as regulators of apoptosis in cancer. Cancer Lett, 2017. 396: p. 53–65.

30. Kearney, C.J., et al., Tumor immune evasion arises through loss of TNF sensitivity. Sci Immunol, 2018. 3(23).

31. Heard, A., et al., Antigen glycosylation regulates efficacy of CAR T cells targeting CD19. Nat Commun, 2022. 13(1): p. 3367.

32. Jongsma, M.L.M., et al., The SPPL3-defined glycosphingolipid repertoire orchestrates HLA class I-mediated immune responses. Immunity, 2021. 54(2): p. 387.

33. Joung, J., et al., CRISPR activation screen identifies BCL-2 proteins and B3GNT2 as drivers of cancer resistance to T cell-mediated cytotoxicity. Nat Commun, 2022. 13(1): p. 1606.

34. Taniguchi, N. and Y. Kizuka, Glycans and cancer: role of N-glycans in cancer biomarker, progression and metastasis, and therapeutics. Adv Cancer Res, 2015. 126: p. 11–51.

35. Lau, K.S. and J.W. Dennis, N-Glycans in cancer progression. Glycobiology, 2008. 18(10): 1. p. 750–60.

36. Hugonnet, M., et al., The Distinct Roles of Sialyltransferases in Cancer Biology and Onco-Immunology. Front Immunol, 2021. 12: p. 799861.

37. de-Souza-Ferreira, M., E.E. Ferreira, and J.C.M. de-Freitas-Junior, Aberrant N-glycosylation in cancer: MGAT5 and beta1,6-GlcNAc branched N-glycans as critical regulators of tumor development and progression. Cell Oncol (Dordr), 2023.

38. Greco, B., et al., Disrupting N-glycan expression on tumor cells boosts chimeric antigen receptor T cell efficacy against solid malignancies. Sci Transl Med, 2022. 14(628): p. eabg3072.

39. Nakano, M., et al., Bisecting GlcNAc Is a General Suppressor of Terminal Modification of N-glycan. Mol Cell Proteomics, 2019. 18(10): p. 2044–2057.

40. Chiba, M., et al., Genome-wide CRISPR screens identify CD48 defining susceptibility to NK cytotoxicity in peripheral T-cell lymphomas. Blood, 2022. 140(18): p. 1951–1963.

41. Joung, J., et al., Genome-scale CRISPR-Cas9 knockout and transcriptional activation screening. Nat Protoc, 2017. 12(4): p. 828–863.

42. Li, W., et al., Quality control, modeling, and visualization of CRISPR screens with MAGeCK-VISPR. Genome Biol, 2015. 16: p. 281.

43. Hobohm, L., et al., N-terminome analyses underscore the prevalence of SPPL3-mediated intramembrane proteolysis among Golgi-resident enzymes and its role in Golgi enzyme secretion. Cell Mol Life Sci, 2022. 79(3): p. 185.

44. Fang, P., et al., A streamlined pipeline for multiplexed quantitative site-specific N-glycoproteomics. Nat Commun, 2020. 11(1): p. 5268.

45. Perez-Riverol, Y., et al., The PRIDE database resources in 2022: a hub for mass spectrometry-based proteomics evidences. Nucleic Acids Res, 2022. 50(D1): p. D543–D552.

